# The HPA stress axis shapes aging rates in long-lived, social mole-rats

**DOI:** 10.1101/2020.02.22.961011

**Authors:** Arne Sahm, Steve Hoffmann, Philipp Koch, Yoshiyuki Henning, Martin Bens, Marco Groth, Hynek Burda, Sabine Begall, Saskia Ting, Moritz Goetz, Paul Van Daele, Magdalena Staniszewska, Jasmin Klose, Pedro Fragoso Costa, Matthias Platzer, Karol Szafranski, Philip Dammann

## Abstract

Sexual activity and/or reproduction doubles life expectancy in the long-lived rodent genus *Fukomys*. To investigate the molecular mechanisms underlying this phenomenon, we analyzed a total of 636 RNA-seq samples across 15 tissues. This analysis suggests that the differences in life expectancy between reproductive and non-reproductive mole-rats are mainly caused by critical changes in the regulation of the hypothalamic-pituitary-adrenal stress axis, which we further substantiate with a series of independent evidence. In accordance with previous studies, the up-regulation of the proteasome and several so-called “anti-aging molecules”, such as DHEA, is also linked with enhanced life expectancy. On the other hand, several our results oppose crucial findings in short-lived model organisms. For example, we found the up-regulation of the IGF1/GH axis and several other anabolic processes to be compatible with a considerable lifespan prolongation. These contradictions question the extent to which findings from short-lived species can be transferred to longer-lived ones.

## Introduction

Most of our current understanding of the underlying mechanisms of aging comes from short-lived model species. It is, however, still largely unclear to what extent insights obtained from short-lived organisms can be transferred to long-lived species, such as humans (Parker et al. 2004; Keller and Jemielity 2006). Comparative approaches, involving species with particularly long healthy lives and seeking the causative mechanisms that distinguish them from shorter-lived relatives try to overcome this limitation (Austad 2009). Many studies that involved organisms with particularly long lifespans, e.g., queens in social hymenoptera, birds, bats, African mole-rats, and primates, have produced findings that were not always congruent with established aging theories (Keller and Jemielity 2006; Austad 2009; Salmon et al. 2009; Austad 2011; Dammann 2017; Bens et al. 2018).

Species comparisons, however, also have their limitations. Many observed differences between species with differing lifespans are influenced by phylogenetic constraints, eco-physiological differences, or both, rather than being causal for the species-specific differences in aging rate and longevity. Therefore, the results mentioned above would gain much more value if they could be confirmed or falsified in a system in which it is unnecessary to cross the species border, that is, in a species that contains easily distinguishable cohorts with high vs. low longevity.

Bimodal aging occurs naturally in the genus *Fukomys* from the rodent family Bathyergidae (African mole-rats). These animals live in families (often called colonies) of usually consisting of 9 to 16 individuals (Sichilima et al. 2008; Skliba et al. 2012), although single families may occasionally grow considerably larger in some species (Jarvis and Bennett 1993; Scharff et al. 2001). Regardless of group size, an established family typically consists of only one breeding pair (the founders of the family, often called king/queen) and their progeny from multiple litters (often called workers). Because of strict avoidance of incest (Burda 1995), the progeny do not engage in sexual activity in the confines of their natal family, even after reaching sexual maturity. Hence, grown *Fukomys* families are characterized by a subdivision into breeders (the founder pair) and non-breeders (all other family members). Interestingly, breeders reach the age of 20 years or more in captivity, whereas non-breeders usually die before their tenth birth date (Fig. 1A). This divergence of survival probabilities between breeders and non-breeders is found in all *Fukomys* species studied so far, irrespective of sex. Because no difference in diet or workload has been observed between breeders and non-breeders in captivity, status-specific changes of gene expression after the transition from non-breeder to breeder are considered the most likely explanation of the differing lifespans (Dammann and Burda 2006; Dammann et al. 2011).

**Figure 1.**
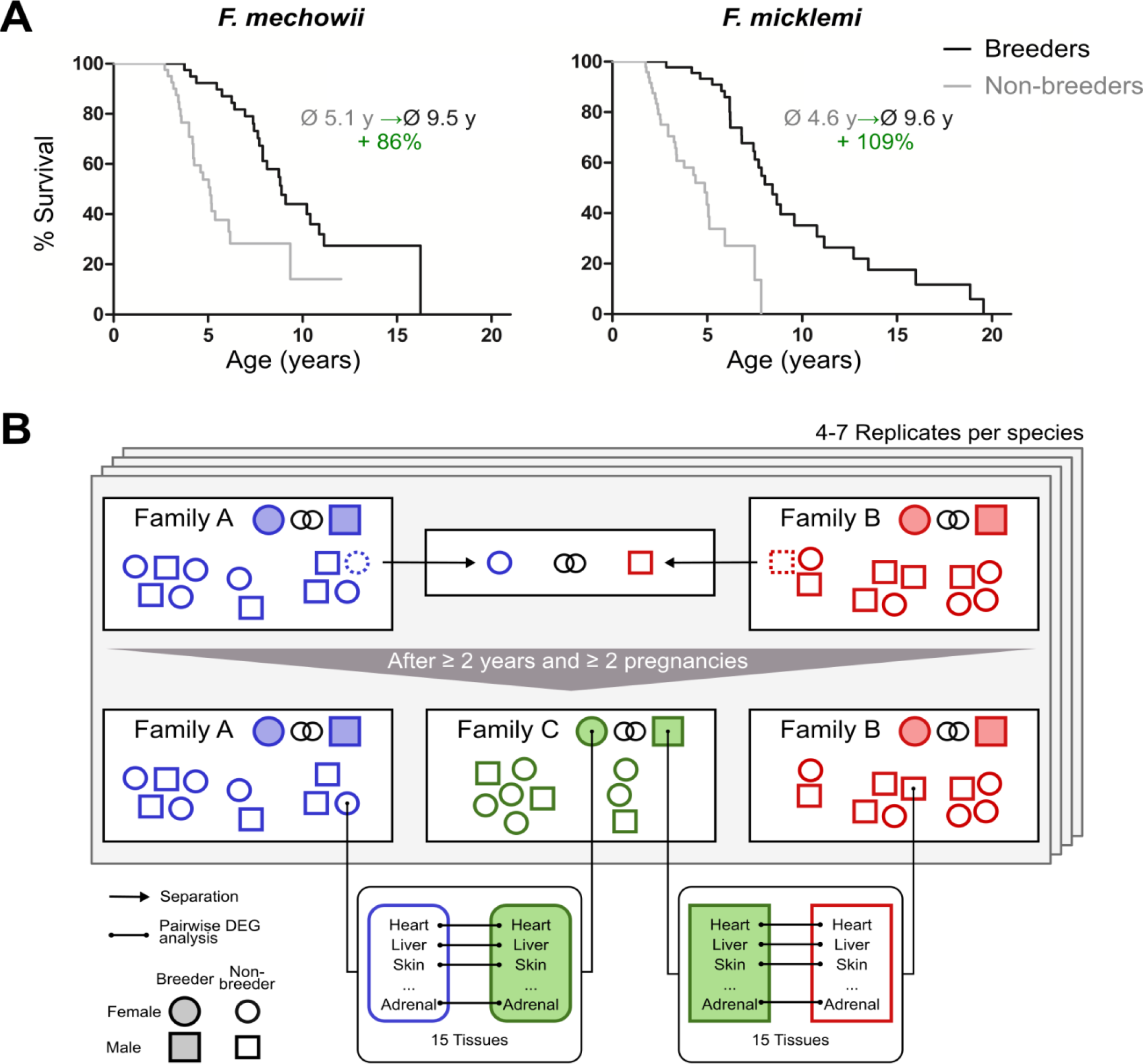
Motivation (A) and principle of the experimental setup (B). **A)** For both species of the *Fukomys* genus that were examined in this study – *F. mechowii* and *F. micklem*i – it was shown that, in captivity, breeder live significantly longer than non-breeders. Lifespan data were redrawn from Dammann & Burda and Dammann et al. (Dammann and Burda 2006; Dammann et al. 2011). **B)** Schematic overview of animal treatments. Non-breeders (open shapes) are offspring of the breeder pair of their family (filled shapes) and do not mate with other members of the same family because of incest avoidance in *Fukomys* (Burda 1995). Therefore, non-breeders of opposite sexes were taken from different families – labeled as “Family A” (blue) and “Family B” (red) – and permanently housed in a separate terrarium. The two unrelated animals mated with each other, thus producing offspring and becoming breeders of the new “Family C” (green). In addition to the animals that were promoted to be slower-aging breeders, age-matched controls that remained in the faster-aging non-breeders of “Family A” and “Family B” were included in our study – in most cases full siblings (ideally litter mates) of the respective new breeders. After at least two years and two pregnancies in “Family C”, breeders from “Family C” and their controls from Colonies A and B were put to death, and tissues were sampled for later analysis, that included identification of differentially expressed genes (DEGs). The shown experimental scheme was conducted with 5 to 7 (median 6) specimens per cohort (defined by breeding status, sex, species) and 12 to 15 tissues (median 14) per specimen: in total, 46 animals and 636 samples.

In the wild, non-breeders must meet a member of another family by chance to ascend to breeder status; in captivity, the establishment of new breeder pairs is subject to human control. Allowing an animal to breed in captivity can be regarded as a simple experimental intervention that results in an extension of life expectancy of approximately 100%. This extension is far more than most experimental interventions in vertebrates can achieve e.g. by caloric restriction (e.g., (Carmona and Michan 2016)) or diets containing resveratrol or rapamycin (e.g., (Valenzano et al. 2006; Johnson et al. 2013)). Furthermore, this relative lifespan extension starts from a non-breeder lifespan that is already more than twice as long as that of the mammalian model organisms most widely used in aging research, such as mice or rats.

Until now, relatively few studies have addressed the potential mechanisms behind this natural status-dependency of aging in *Fukomys* sp. Contrary to the predictions of both the advanced glycation formation theory (Morimoto and Cuervo 2009) and the oxidative stress theory of aging (Harman 1956; Harman 2001), markers of protein cross-linking and -oxidation were surprisingly higher in breeders of Anselĺs mole-rats (*F. anselli*) than in age-matched non-breeders (Dammann et al. 2012). On the other hand, in the Damaraland mole-rat (*F. damarensis*), oxidative damage to proteins and lipids was significantly lower in breeding females than in their non-reproductive counterparts (Schmidt et al. 2014), a finding that is compatible with the oxidative stress theory of aging. In good agreement with their overall longevity irrespective of social status, Anselĺs mole-rats produce less thyroxine (T4) and recruit smaller proportions of their total T4 resources into the active unbound form than do euthyroid mammals. Still, nonetheless the levels of unbound T4 (fT4) do not explain the intraspecific differences in aging rates between *F. anselli* breeders and non-breeders, because the levels of this hormone did not differ between the two cohorts (Henning et al. 2014). Closely connected to the topic of this paper is the finding, that non-breeding giant mole-rats (*F. mechowii*) maintain fairly stable gene expression into relative old ages, quite in contrast to the shorter-lived Norway rat (Sahm et al. 2018a). It is, however, still unclear what happens on the gene expression level when an individual attains breeding status.

Interestingly, a recent study by (Bens et al. 2018) found that the longest-lived rodent, the naked mole-rat *Heterocephalus glaber*, tended globally to show opposite changes in the transition from non-breeders to breeders compared to shorter-lived guinea pigs. *Heterocephalus* and *Fukomys* are similar in their mating and social behavior, but differences appear to exist regarding the effect of breeding on aging rates: until very recently, naked mole-rat non-breeders have been reported to be as long-lived as breeders (Sherman and Jarvis 2002; Buffenstein 2008). In 2018, a lifespan advantage of breeders over non-breeders was reported in females (but not males), yet the divergence of the two groups appeared to be considerably smaller than in *Fukomys* (Ruby et al. 2018) and underlying data is being debated (Dammann et al. 2019). In summary, status-dependent aging is either absent in *Heterocephalus* or less pronounced than in *Fukomys*.

In this paper, we make use of the bimodal aging pattern in two *Fukomys* species (*F. mechowii* and *F. micklemi*, Fig. 1A) by comparing the gene expression profiles of breeders (n=24) and age-matched non-breeders (n=22) in 16 organs or their substructures (hereinafter referred to as tissues, Fig. S1, Fig. 1B). Our main aim was to identify genes and pathways whose transcript levels are linked to the status-dependent aging-rates and to relate these patterns to insights into aging research obtained in shorter-lived species.

## Results

We measured gene expression differences between breeders and non-breeders in two African mole-rat species, *F. mechowii* and *F. micklemi*. Altogether, we performed RNA-seq for 636 tissue samples covering 16 tissue types from both species, sexes, and reproductive states (breeders and non-breeders). Each of the four groups (male/female breeders/non-breeders) of each species consisted of 5 to 7 animals (see Tables S1 –S5 for sample sizes, animal data, and pairing schemes). For each tissue, we conducted a multi-factorial analysis of differentially expressed genes (DEGs): the analysis was based on the variables reproductive state, sex, and species. During this exercise, we focused on the differences between slower-aging breeders and faster-aging non-breeders. This approach increases our statistical power by giving us a four-fold increase of sample size in comparison to species- and sex-specific breeder vs. non-breeder analyses. At the same time, we can additionally reduce the number of false-positive DEGs by restricting the analysis to those breeding status-related genes that show the same direction in both sexes and both species. We deliberately focused on those genes to concentrate our study on universal mechanisms that hold for both sexes and species.

To globally quantify the transcriptomic differences between the reproductive states, we performed three analyses: clustering of the samples based on pairwise correlation, principal variant component analysis, and an overview of the number of DEGs between reproductive states in comparison to DEGs between species and sex. Clustering of the samples based on pairwise correlations showed a full separation of the two species at the highest cluster level (Fig. S1). Below that level, an almost complete separation according to tissues was observed. Within the tissue clusters, the samples did not show a clear-cut separation between sex or breeder/non-breeder status. Accordingly, a principal variance component analysis showed that species, tissue, and the combination of both variables accounted for 98.4 % of the total variance in the data set; individual differences explained 1.4 % of the variance, and only 0.004 % was explained by breeder/non-breeder status (Fig. 2A). Regarding the numbers of DEGs, we found – unsurprisingly considering the aforementioned facts –by far the highest number of DEGs in the species comparison (Fig. 2B). Although in almost every examined tissue the numbers of detected DEGs were also high between sexes, most tissues exhibited very few DEGs due to breeder/non-breeder status. Exceptions were liver, spleen, ovary and, especially, tissues of the endocrine system (adrenal gland, pituitary gland, thyroid), in which the number of DEGs between breeders and non-breeders ranged from more than sixty to several thousand.

**Figure 2.**
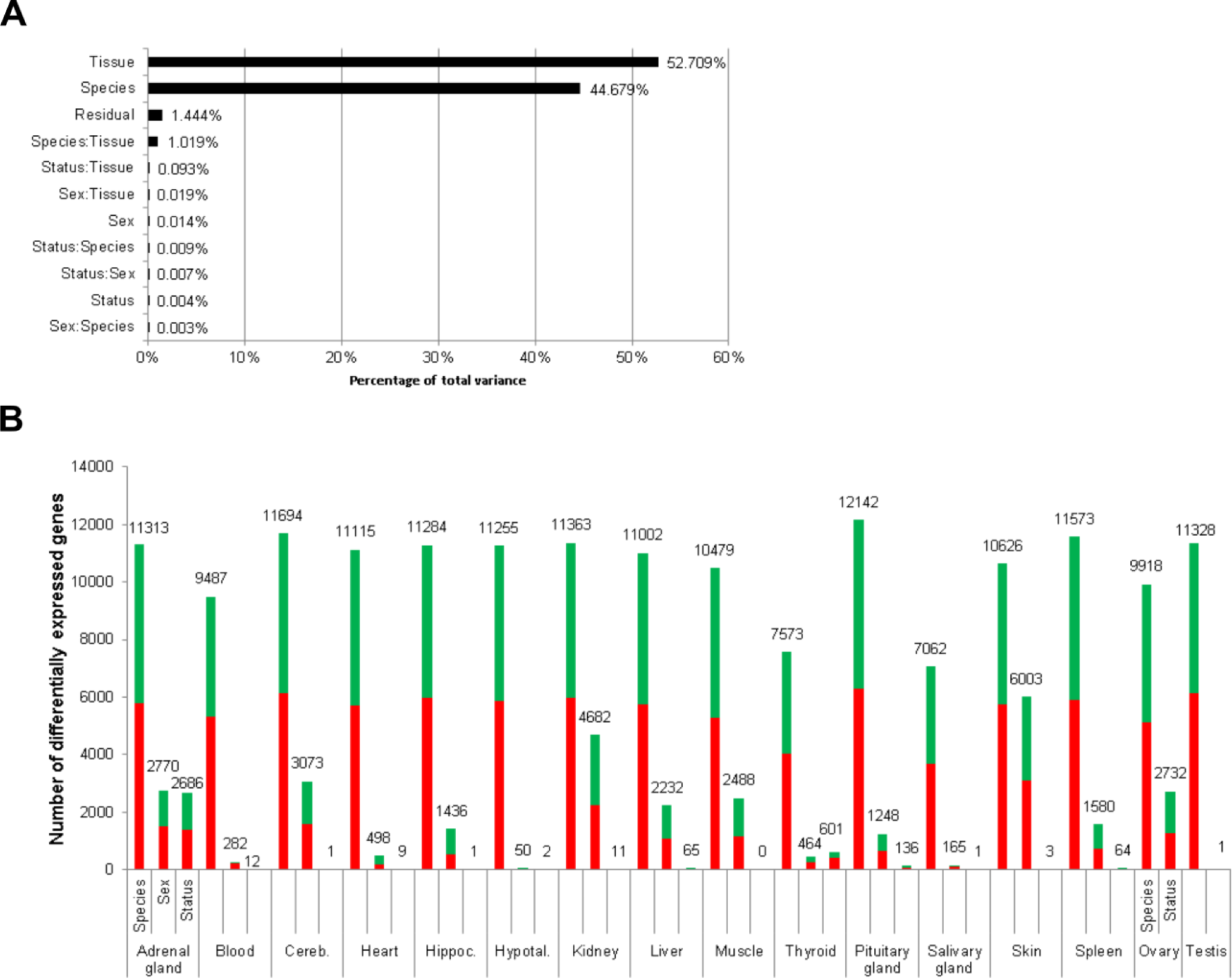
Total variance distribution (A) and numbers of differentially expressed genes (B). **A)** Relative contribution of the model factors (breeding status, sex, species, tissue) and their combinations (:) to the total variance in the examined data set. The relative contributions were determined by principal variance component analysis. **B)** Numbers of identified differentially expressed genes per tissue and model factor (first column, species; second, sex; third, status). Stacked bars indicate the proportions of up- and down-regulated genes (red and green, respectively; directions: *F. mechowii* vs. *F. micklemi*, female vs. male, breeder vs. non-breeder).

Next, we evaluated the relevance of reproductive status DEGs for aging and aging-related diseases. For this analysis, we first determined overlaps by using the Digital Aging Atlas (DAA) – a database of genes that show aging-related changes in humans (Craig et al. 2015). Across species and sexes, significant overlaps (FDR < 0.05, Fisher’s exact test) with the DAA were found in three tissues: adrenal gland, ovary and pituitary gland (false discovery rate [FDR] = 0.005, each; Fig. 3A). Among these three endocrine tissues, the DEGs of the ovaries overlapped significantly with those from adrenal (p=2.8*10^-27^) and pituitary glands (p=0.005), but there was no significant overlap between the two glands (Fig. 3A). Thus, together, we found indications for aging-relevant expression changes after the transition from non-breeders to breeders in three tissues of the endocrine system, which presumably affect separate aspects of aging in adrenal and pituitary glands.

Moreover, we compared the DEGs with respect to the reproductive status that we identified in *Fukomys* with regard to their direction to transcript-level changes observed in similar experiments using naked mole-rats and guinea pigs (Bens et al. 2018). The direction of the status-dependent DEGs regulation in *Fukomys*, as found in this study, was significantly more often the same rather than opposite as in the naked mole-rat (females, 60 %, p=5.2*10^-58^; males, 62 %, p=10^-44^ for females, Fig.3B, Table S6). In the guinea pig, on the contrary, the *Fukomys* reproductive status DEGs were significantly more often regulated in the opposite direction (females, 57 %, p=9*10^-25^; males, 59 %, p=4.7*10^-23^, Fig. 3B, Table S6). Thus, at the single-gene level, the expression changes linked to reproductive status may affect lifespan differently in long-lived African mole-rats than in shorter-lived guinea pigs.

Beyond the single-gene level, we aimed to identify metabolic pathways and biological functions whose gene expression significantly depends on reproductive status. For this, we used Kyoto Encyclopedia of Genes and Genomes (KEGG) pathways (Kanehisa et al. 2017), and Molecular Signatures Database (MSigDB) hallmarks (Liberzon et al. 2015) as concise knowledge bases. We used a known method that combines all p-values of genes in a given pathway in a threshold-free manner. The advantage of this approach is that it bundles the p-values from test results of individual gene expression differences at the level of pathways (see Methods section for details). Altogether, the gene expression of 55 KEGG pathways and 41 MSigDB hallmarks was significantly affected by reproductive status in at least one tissue (Fig. S3 and S4). Because the individual interpretation of each of these pathways/hallmarks would go beyond the scope of this study, we focus here on those 14 pathways and 13 hallmarks that were significantly different between breeders and non-breeders (FDR < 0.1) in a global analysis across all tissues (Fig. 4). Because many pathways are driven mainly by gene expression in subsets of tissues, we weighted gene-wise the differential expression signals from the various tissues by the respective expression levels in the tissues. For instance, the expression level of the growth hormone (GH) gene *GH1* is known to be almost exclusively expressed in the pituitary glands. In our data set the *GH1* level of the pituitary gland accounted for 99.96 % of the total *GH1* across all tissues. Accordingly, in pathways that contain *GH,* our weighted cross-tissue differential expression signal for this gene is almost exclusively determined by the pituitary gland. On the contrary, a differential expression signal of this gene in another tissue with a very low fraction of the gene’s total expression would have almost no impact on the weighted cross-tissue level – even if that signal were very strong (see Methods section for details).

**Figure 3.**
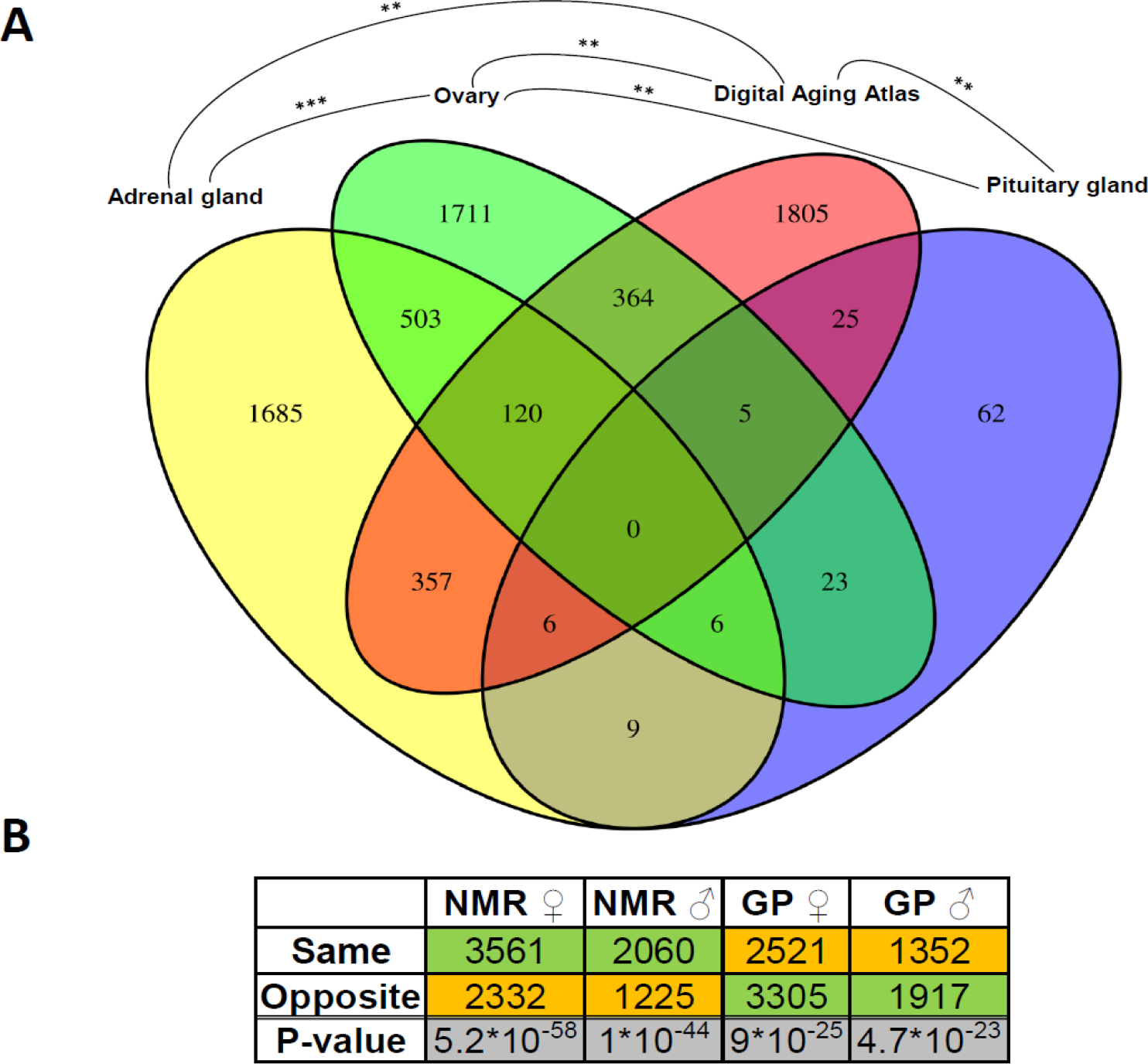
Assessment of the aging relevance of genes that are differentially expressed between breeders and non-breeders. **A)** For each tissue, we separately tested whether the identified differentially expressed genes between status groups significantly overlapped with the genes within the Digital Aging Atlas database (Fisher’s exact test, FDR < 0.05). Significant overlaps were found for three tissues: adrenal gland, ovary and pituitary gland. The Venn-diagram depicts the overlaps of these three tissues with the Digital Aging Atlas and with each other (**: FDR < 0.01; ***: FDR < 0.001). **B)** A similar experiment comparing the transcriptomes of breeders versus non-breeders was recently conducted in naked mole-rats (NMRs) and guinea pig (GPs) (Bens et al. 2018). For NMR there is also evidence that breeders have a (slightly) longer lifespan than non-breeders, whereas for GP the opposite is assumed (Bens et al. 2018; Ruby et al. 2018). Across ten tissues that were examined in both studies, the analysis determined whether status-dependent differentially expressed genes identified in the current study were regulated in the same or opposite direction in NMR and GP (Table S6). The listed p-values (two-sided binomial exact test; hypothesized probability, 0.5) describe how extremely the ratio of genes expressed in the same and opposite directions deviates from a 50:50 ratio. Green and orange indicate the majority and minority of genes within a comparison, respectively. Figure created with BioRender.com.

**Figure 4.**
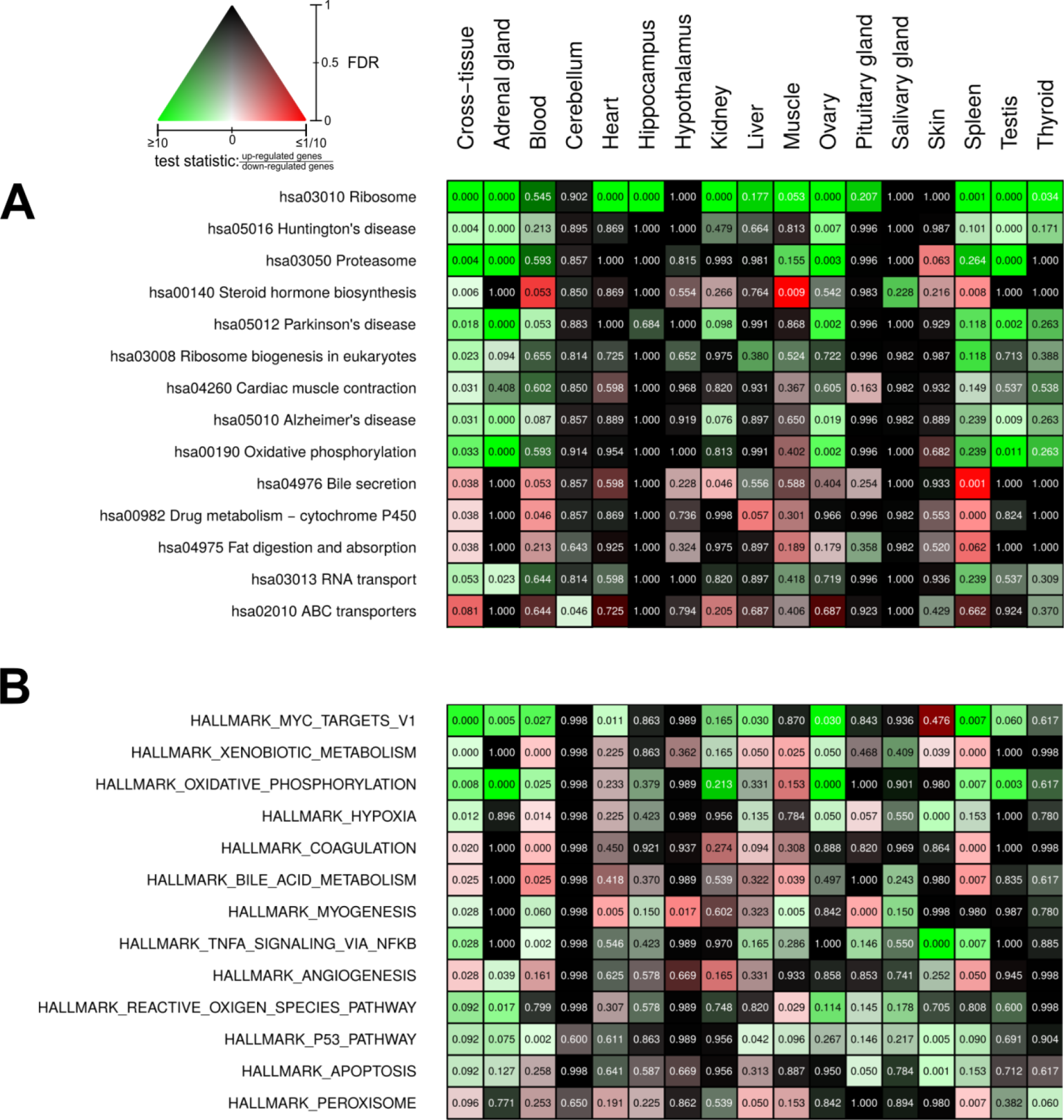
Pathways and metabolic functions enriched for status-dependent differential gene expression. Shown are all Kyoto Encyclopedia of Genes and Genomes (KEGG) pathways (**A**) and Molecular Signatures Database (MSigDB) hallmarks (**B**) that are enriched for differential gene expression between breeders and non-breeders at the weighted cross-tissue level (false discovery rate [FDR], < 0.1). The numbers within the cells give the FDR, i.e., the multiple testing corrected p-value, for the respective pathway/hallmark and tissue. As indicated by the color key, red and green stand for up- or down-regulated in breeders, respectively. White indicates a pathway/hallmark that is significantly affected by differential expression and whose signals for up- and down-regulation are approximately balanced. Dark colors up to black mean that there is little or no evidence that the corresponding pathway/hallmark is affected by differential gene expression. Figures S3 and S4 provide detailed overviews of all pathways/hallmarks that are enriched in at least one tissue for status-dependent differential expression signals.

We found strong indications for increase in the activity of certain anabolic functions in breeders: Ribosomal protein expression (hsa03010 Ribosome, hsa03008 Ribosome biogenesis in eukaryotes, Fig. 4A) was elevated in most tissues (and accordingly also in the weighted cross-tissue analysis). In MSigDB hallmarks, the strongest enrichment signal came from MYC targets (HALLMARK_MYC_TARGETS_V1), which can largely be considered a reflection of enhanced ribosomal protein expression and the fact that MYC is a basal transcription factor up-regulating genes involved in protein translation ((Hofmann et al. 2015), Fig. 4B). In functional correspondence, we observed an increase in the expression of mitochondrial respiratory chain components (hsa00190 Oxidative Phosphorylation, HALLMARK_OXIDATIVE_PHOSPHORYLATION, Fig. 4A,B). We also found strong indications for increased protein degradation (hsa03050 PROTEASOME, Fig. 4A). This weighted cross-tissue signal was, in contrast to the situation regarding ribosomes, driven mainly by two tissue types: the adrenal gland and the gonads.

To examine whether the simultaneous up-regulation of the ribosome, proteasome, and oxidative phosphorylation is a coordinated regulation, we performed a weighted gene co-expression network analysis (WGCNA) (Langfelder and Horvath 2008) from our gene count data and examined the connectivity between pairs of those KEGG pathways flagged in the weighted cross-tissue analysis. We found that the expression of ribosomal genes (hsa03010) was significantly linked to those of ribosome biogenesis (hsa03008, FDR=4.59*10^-3^), oxidative phosphorylation (hsa00190, FDR=4.05*10^-4^), and proteasome (hsa03050, FDR=4.59*10^-3^), whereas no other examined pathway pair exhibited a significant connectivity (Fig. S5). Interestingly, ribosome, proteasome, and oxidative phosphorylation pathways also shared other characteristics of their differential expression signals: subtle fold-changes, that is, up-regulation of 3 to 9 % on average. Thus, statistically significant signals at the pathway level resulted from relatively small shifts in all genes of these pathways in a seemingly coordinated manner and across multiple tissues (Data S1). In addition, ribosome (hsa03010, in ovary), proteasome (hsa03050, in ovary and adrenal gland), and RNA-transport (hsa03013, in adrenal gland) are enriched in those *Fukomys* status-dependent DEGs that show, in a similar experimental setting (Bens et al. 2018), the same direction in both naked mole-rat sexes and the opposite direction in both guinea pig sexes (FDR < 0.05, Fisher’s exact test).

The myogenesis hallmark (Fig. 4B) was also found to be up-regulated in breeders. Expectedly, this weighted cross-tissue result was driven mainly by differential expression signals from muscle tissue: muscle from all tissues exhibited the lowest p-value (Fig. 4A), and 15 of 20 up-regulated genes that contributed most to the weighted cross-tissue differential myogenesis signal exhibited their highest expression in muscle. These genes were involved mainly in calcium transport or part of the fast-skeletal muscle-troponin complex (Data S1). A clear exception of this muscle-dominated expression is found in the gene that exhibited the highest relative contribution to the differential myogenesis signal, *insulin-like growth factor 1* (*IGF1*). This gene was found to be expressed most strongly in ovary and liver and was strongly up-regulated in the breederś ovaries and adrenal glands (Table 1). *IGF1* codes for a well-known key regulator of anabolic effects such as cell proliferation, myogenesis, and protein synthesis (Schiaffino and Mammucari 2011; Jung and Suh 2014) and has a tight functional relation to GH (gene: *GH1*) another key anabolic regulator upstream of *IGF1*; together, these factors form the so-called GH/IGF1 axis (Cannata et al. 2010; Junnila et al. 2013; Bodart et al. 2015; Raisingani et al. 2017; Carotti et al. 2018; Lozier et al. 2018). Also, *GH1* was strongly up-regulated in breeders in its known principal place of synthesis, the pituitary gland (Table 1).

**Table 1.** Top ten genes regarding weighted cross-tissue differential expression signal.

|  | Weighted cross-tissue |  |  | Tissue with highest expression |  |  |  |
| --- | --- | --- | --- | --- | --- | --- | --- |
|  | P-value | FDR | Log2-foldchange* | Tissue | % of cross-tissue Expression | FDR | Log2-foldchange* |
| **SULT2A1** | 0.00E+00 | 0.000 | -3.19 | Liver | 97.42 | 5.27E-06 | -3.26 |
| **MC2R** | 6.00E-06 | 0.046 | -0.53 | Adrenal gland | 89.53 | 8.78E-05 | -0.56 |
| **INHA** | 1.60E-05 | 0.081 | 1.57 | Ovary | 67.91 | 1.54E-05 | 2.22 |
| INHA – tissue with 2<sup>nd</sup> highest expression |  |  |  | Testis | 27.61 | 0.56 | 0.21 |
| **CYP11A1** | 5.90E-05 | 0.224 | 0.61 | Ovary | 45.55 | 5.49E-05 | 0.91 |
| CYP11A1 – tissue with 2<sup>nd</sup> highest expression |  |  |  | Adrenal gland | 43.63 | 1.48E-03 | 0.47 |
| **NLRP14** | 1.17E-04 | 0.355 | -0.80 | Ovary | 68.98 | 2.84E-05 | -1.25 |
| NLRP14 – tissue with 2<sup>nd</sup> highest expression |  |  |  | Testis | 18.67 | 0.54 | 0.19 |
| **GH** | 2.58E-04 | 0.618 | 0.45 | Pituitary gland | 99.96 | 1.99E-02 | 0.45 |
| **IGF1** | 3.62E-04 | 0.618 | 0.59 | Ovary | 51.73 | 1.20E-04 | 0.76 |
| IGF1 – tissue with 2<sup>nd</sup> highest expression |  |  |  | Liver | 36.28 | 0.39 | 0.24 |
| **ZP4** | 3.74E-04 | 0.618 | -1.16 | Ovary | 95.30 | 1.99E-03 | -1.23 |
| **TCL1A** | 5.33E-04 | 0.618 | -1.05 | Ovary | 90.12 | 2.14E-03 | -1.18 |
| **PNLIPRP2** | 5.69E-04 | 0.618 | 1.36 | Ovary | 76.08 | 2.23E-03 | 1.65 |

\* Direction: breeder/non-breeder
Note – tissues are listed if they contribute at least 10% to cross-tissue expression

With xenobiotic metabolism and TNF-α-signaling, two defense hallmarks were also found to be up-regulated in breeders by the weighted cross-tissue analysis (HALLMARK_XENOBIOTIC_METABLISM and HALLMARK_TNFA_SIGNALING_VIA_NFKB, Fig. 4B). The up-regulation of the reactive oxygen species (ROS) hallmark comprising genes coding for proteins that detoxify ROS (HALLMARK_REACTIVE_OXYGEN_SPECIES_PATHWAY, Fig. 4B) falls into a similar category.

Another interesting aspect that was found to be significantly altered in breeders is steroid hormone biosynthesis (hsa00140 Steroid hormone biosynthesis, Fig. 4A). In this case, both up- and down-regulated genes were in the pathway, and their absolute fold-changes were roughly balanced. Steroid hormones, on the one hand, comprise sex steroids – because these hormones are important players in sexual reproduction, such differences should be expected given the experimental setup. On the other hand, the class of steroid hormones – corticosteroids – has regulatory functions in metabolism, growth, and the cardiovascular system, as well as in the calibration of the immune system and response to stress (Liu et al. 2013). The most influential contributor to the differential pathway signal by far on the weighted cross-tissue level was *CYP11A1,* which codes for the (single) enzyme that converts cholesterol to pregnenolone. This is the first and rate-limiting step in steroid hormone synthesis (Miller and Auchus 2011). Because *CYP11A1* was found to be up-regulated in breeders in its main places of synthesis – the gonads and the adrenal gland (Table 1) – it can be assumed that the total output of steroid hormone biosynthesis in breeders is increased. The pattern of up- and down-regulation on the KEGG pathway, however, suggests that sex steroids especially are produced at a higher rate in breeders whereas circulating levels of glucocorticoids – such as cortisol – should be lower than in non-breeders (Fig. S6; see also our discussion on ACTH-R below).

Finally, several pathways flagged by the weighted cross-tissue analysis seem to be derivatives of the above-mentioned differentially expressed pathways instead of representing altered functions on their own. For example, Huntington’s (hsa05016), Parkinson’s (hsa05012), and Alzheimer’s (hsa05010) diseases could, in principle, be interpreted as highly relevant for aging and lifespan. A closer inspection of these pathways reveals, however, that the genes of the mitochondrial respiratory chain – which is the core of the oxidative phosphorylation pathway – are in all three cases the main contributors to the respective differential expression signals (Fig. S7-S9, Data S1). Similarly, we see in the case of fat digestion (hsa04975 Fat digestion and absorption) that two of the three largest contributors to the differential expression signal of that pathway – *ABCG8* and *SCARB1 –* are directly involved in the transport of cholesterol (Liu et al. 2008; Wang et al. 2015). Therefore, it seems likely that this signal is an expression of the altered steroid hormone biosynthesis rather than indicating altered fat digestion.

Interestingly, three genes, which we had already mentioned as potential regulators during the pathway analysis, also appeared among the ten most clearly altered genes on the weighted cross-tissue level: *GH1, IGF1,* and *CYP11A1* (Table 1). The top two among these ten are *sulfotransferase family 2A member 1* (*SULT2A1*) and *melanocortin 2 receptor* (*MC2R*). SULT2A1 is the main catalyzer of the sulfonation of the steroid hormone dehydroepiandrosterone (DHEA) to its non-active form DHEA-S (Hammer et al. 2005). DHEA has repeatedly been proposed to be an “anti-aging hormone” because its levels are negatively associated with chronological aging (Baulieu 1996; Celec and Starka 2003; Rutkowski et al. 2014). We found that *SULT2A1* is strongly down-regulated in breeders’, liver which is also the main location of its enzymatic action. The second candidate, *MC2R*, encodes the adrenocorticotropin (ACTH)-receptor, which is the main inducer of glucocorticoid synthesis and a crucial component of the hypothalamic–pituitary–adrenal (HPA) axis (Walker et al. 2015). In humans and many other mammals, prolonged glucocorticoid excess leads to Cushing’s syndrome. Affected individuals exhibit muscle weakness, immune suppression, impairment of the GH/IGF1 axis, higher risk of diabetes, cardiovascular disease (hypertension), osteoporosis, decreased fertility, depression, and weight gain (Chabre 2014; Ferrau and Korbonits 2015). The large overlap of these symptoms with those of aging could explain to some extent that Cushing’s syndrome patients exhibit considerably higher mortality rates (Etxabe and Vazquez 1994). We hypothesized that the increased expression of the ACTH receptor in *Fukomys* non-breeders can cause similar expression patterns and consequences (Fig. 5).

**Figure 5.**
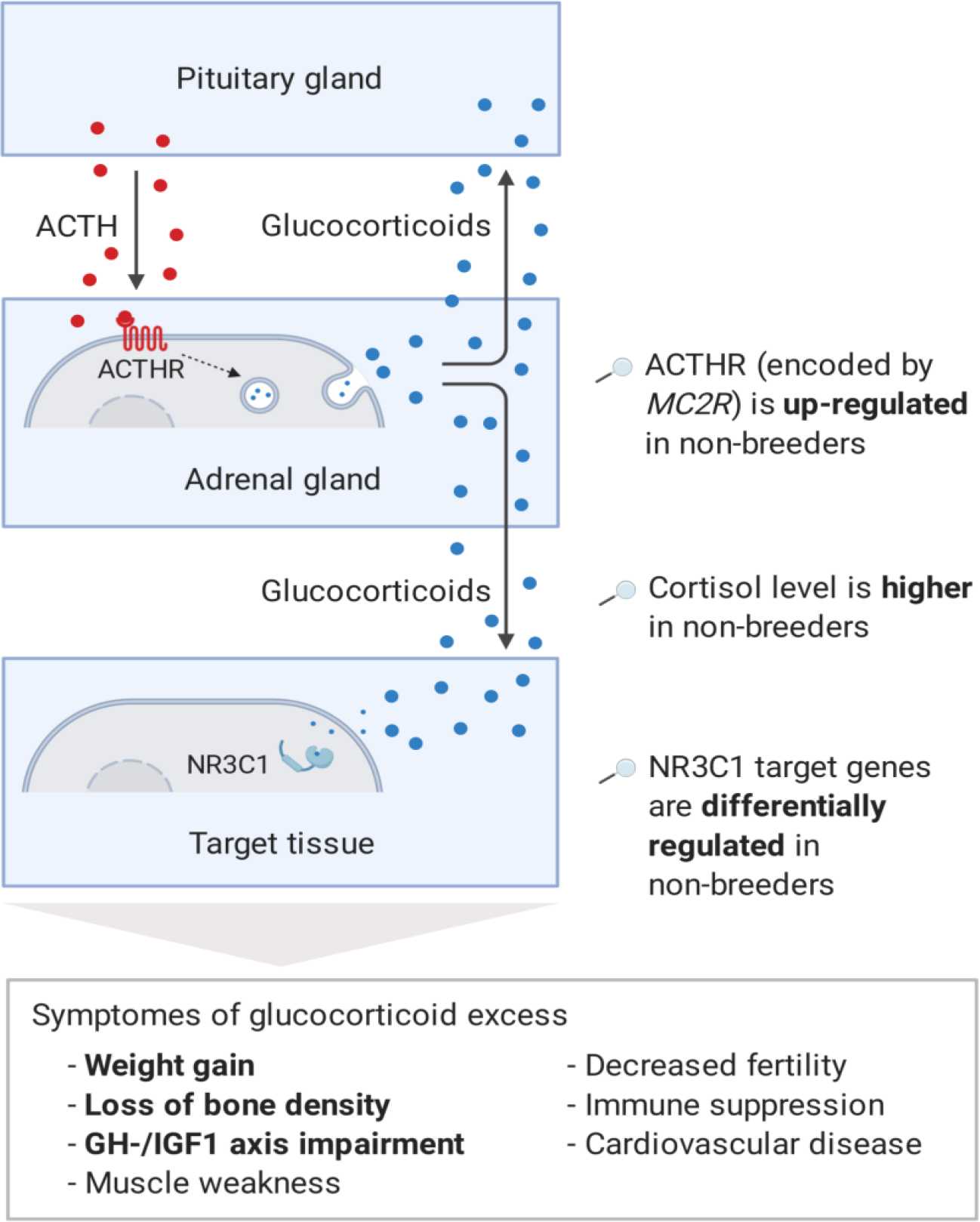
Model of the stress axis as a key mechanism for status-dependent lifespan differences in *Fukomys*. From a wide range of mammals, including humans (Ferrau and Korbonits 2015), dogs (Kooistra and Galac 2012), horses (McCue 2002), cats (Meijer et al. 1978), and guinea pigs (Zeugswetter et al. 2007), it is known that chronic glucocorticoid excess leads to a number of pathologic symptoms that largely overlap with those of aging and result in considerably higher mortality rates for affected individuals (Etxabe and Vazquez 1994#808). The most common cause of chronic glucocorticoid excess is excessive secretion of the adrenocorticotropic hormone (ACTH) by the pituitary gland. ACTH is transported via the blood to the adrenal cortex where it binds to the ACTH-receptor (ACTHR; encoded by the gene *MC2R*) which induces the production of glucocorticoids, especially cortisol. Glucocorticoids are transported to the various tissues, where they exert their effect by activating the glucocorticoid receptor (NR3C1) that acts as a transcription factor and regulates hundreds of genes. The constant overuse of this transcriptional pattern eventually leads to the listed symptoms. Our hypothesis is that the permanent, high expression of the ACTH-receptor in *Fukomys* non-breeders causes effects similar to those known from overproduction of the hormone. In line with this hypothesis, i) cortisol levels are increased in non-breeders and ii) target genes of the glucocorticoid-receptor are highly enriched for status-dependent differential gene expression. Furthermore, the animals were examined for common symptoms of chronic glucocorticoid excess: iii) non-breeders gained more weight during the experiment than breeders, iv) exhibited lower bone density at the end of the experiment, and v) lower gene expression in the GH-/IGF1 axis than breeders.

We tested this hypothesis (Fig. 5) by checking five of its key predictions. Altered *MC2R* expression (Fig. 6A) coincides with higher cortisol levels in hair samples from non-breeding *F. mechowii* than in those from breeders of the same species (Begall et al., in prep.). Furthermore, glucocorticoids such as cortisol exert their effect by binding to the glucocorticoid receptor that, in turn, acts as a transcription factor for many genes (Gjerstad et al. 2018). We tested whether the expression of targets of the glucocorticoid-receptor (NR3C1) was significantly altered throughout our data using two gene lists (Phuc Le et al. 2005): about three hundred direct target genes of the receptor that were identified by chromatinimmunoprecipitation (i), and about 1300 genes that were found to be differentially expressed depending on the presence or absence of exogenous glucocorticoid (ii). Both gene lists were found to be significantly affected by differential expression at the weighted cross-tissue level (I, p=0.001; ii, p<10^-9^) as well as in 5 (i) and 8 (ii) single tissues (Table S7, S8). In line with our hypothesis, we observed that the weight gain in non-breeders was, on average, twice as strong compared to the weight gain in breeders during the experiment (p=7.49*10^-3^,type II ANOVA, Fig. 6B). In addition, we found a subtle but significant influence of reproductive status on the density of the vertebrae’: the vertebrae of breeders were slightly denser than those of age-matched non-breeders (p=0.03 for vertebra T12 only, and p=0.01 across all examined vertebrae L1, L2, and T12; ANOVA, Fig. 6C, Table S11).

**Figure 6.**
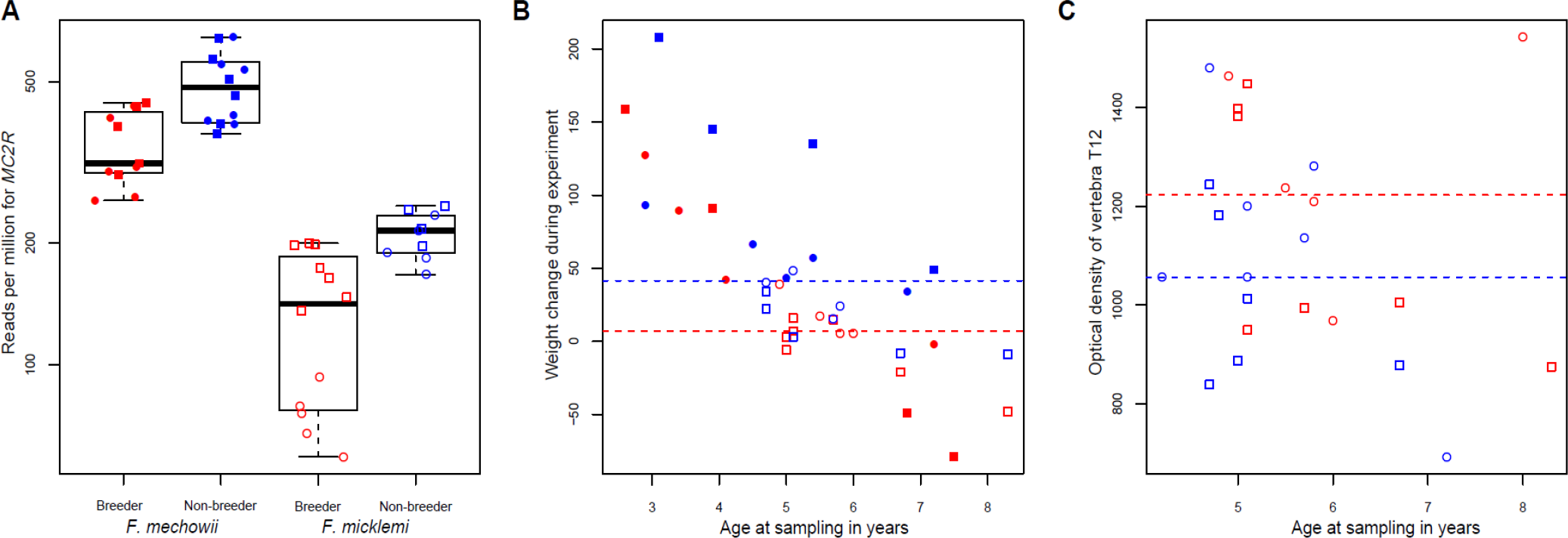
*MC2R* gene expression and physiological measurements. **A)** Gene expression of *MC2R*, coding for the ACTH receptor, in breeders and non-breeders of *Fukomys mechowii* and *Fukomys micklemi*. **B)** Weight gain of the animals during the experiment. **C)** Measured optical densities of vertebra T12 of *Fukomys micklemi* breeders and non-breeders. Red, breeders; blue, non-breeders; filled, *F. mechowii*; unfilled, *F. micklemi*; circles, females; squares, males; dashed line, median. Statistically significant differences between breeders and non-breeders were determined with A) DESeq2 (Love et al. 2014) and B+C) analysis of variance with status, species, sex, and age as independent variables (see methods).

## Discussion

The vast lifespan differences between *Fukomys* breeders and non-breeders are, according to our RNA-seq data, associated with only subtle global pattern shifts in transcript levels. Concerning the tested explanatory variables (*Fukomys* species, sex, breeding status), we found by far the highest number of DEGs at the level of the species comparison. Although the number of DEGs between the sexes was comparably high in almost each examined tissue, only very few DEGs were found comparing breeders and non-breeders. One exception is the ovary, whose high number of DEGs corresponds well with the disparity in reproductive activity. Other exceptions are liver, spleen, and, especially, the tissues of the endocrine system (adrenal gland, pituitary gland, thyroid), in which the number of DEGs between breeders and non-breeders ranged from more than 100 to more than 2,500.

Changes in the gene expression of the endocrine system are well known to play an important role both in sexual maturation an in aging and the development of aging-associated diseases, e.g., diabetes and cardiovascular diseases (Tatar et al. 2003; Chahal and Drake 2007; Jones and Boelaert 2015). This finding fits well with the observation of substantial changes in the endocrine system after the transition from non-breeders to breeders in the related naked mole-rat – one of the key results of a recent study in that species (Bens et al. 2018). The dominance of differential expression in endocrine tissue is also plausible insofar as these tissues exert a strong control function for other tissues via hormone release.

Steroid hormone biosynthesis exhibits a bipartite pattern in breeders, with up-regulated sex steroid genes and a simultaneous down-regulation of corticosteroid synthesis genes. The former could be expected as a consequence of sexual activity in breeders. For aging, it is, however, interesting that *SULT2A1*, a gene that codes for the specialized sulfotransferase converting the sex steroid DHEA to its non-active form DHEA-S was found to be heavily down-regulated in breeders. DHEA is the most abundant steroid hormone; it serves as a precursor for sex steroid biosynthesis but also has various metabolic functions on its own (Allolio and Arlt 2002; Webb et al. 2006). DHEA levels decrease continuously during the human aging process to an extent that favors it as an aging biomarker (Maggio et al. 2007; Traish et al. 2011). As a result, an aggressive advertising of DHEA as “anti-aging hormone” in the form of dietary supplements could be observed in recent years (Baulieu 1996; Celec and Starka 2003; Webb et al. 2006; Rutkowski et al. 2014). Despite conflicting experimental data from various animal studies and clinical trials, a positive effect of DHEA on human health is frequently considered to be likely in the literature. Large-scale and long-term studies, however, are still pending (Traish et al. 2011; Rutkowski et al. 2014; Samaras et al. 2015). Interestingly, DHEA is not only described as an aging marker but also as a marker for clinically relevant glucocorticoid excess (Burkhardt et al. 2013).

Regarding corticosteroid synthesis, we hypothesize that the regulation of adrenal gland *MC2R*, coding for the ACTH receptor, is likely to cause a considerable proportion of the overall observed expression patterns and of the lifespan extension in breeders (Fig. 5). As a critical component of the HPA axis, ACTH is a stress hormone that is produced by the pituitary gland and transported by the the blood to the adrenal cortex, where it binds to the ACTH receptor (Fridmanis et al. 2017). Subsequently, the ACTH receptor induces the synthesis of glucocorticoids (Walker et al. 2015), e.g., cortisol, which in turn cause immunosuppressive and various metabolic effects throughout the organism (Becker 2013). Although in many mammals (such as humans, dogs, and guinea pigs) glucocorticoid excess disorder, Cushing’s syndrome, is caused by overproduction of the hormone ACTH. Our results for *Fukomys* suggest, that the increased levels of the ACTH receptor may lead to symptoms and expression patterns that could resemble some of molecular and phenotypic aspects of this pathological condition (Fig. 5). Our hypothesis is supported by a number of confirmed downstream effects: the target genes of the glucocorticoid receptor are highly significantly affected by differential gene expression; non-breeder symptoms were similar to those of humans with abnormally elevated glucocorticoid levels: weight gain, decreases in bone density, and impairment of the GH/IGF1 axis (Fig. 5, Fig.6). Interestingly, blocking of the ACTH receptor has been suggested as a treatment for human Cushing’s syndrome (Newfield 2010).

We looked for evidence of a regulation upstream of the ACTH receptor. Given the known positive feedback loop between the ACTH receptor and its own ligand, decreased ACTH synthesis in breeders would have been an obvious explanation (Imai et al. 2001). We found that the expression of the ACTH polypeptide precursor gene (*POMC*) is unchanged. However, also other post-transcriptional or post-translational mechanisms such as cleaving may influence the ACTH levels in breeders and non-breeders. Of those genes known to be involved in the regulation of *MC2R* (Beuschlein et al. 2001; Lin et al. 2007), we found that only *PRKAR1B* was differentially regulated in the adrenal gland. However, this gene codes for only one of 7 subunits of the involved protein kinase A. Alternative explanations could be that epigenetic modifications, other still unidentified regulators of the transcript level, or both, are responsible for the differential expression.

In the circulatory system, represented by blood and spleen, down-regulation of coagulation factors was observed in breeders. Because coagulation factors are known to be up-regulated during aging in humans mice, rats, and even fish, this can be interpreted as a sign of a more juvenile breeders’ transcriptome (Ochi et al. 2016; Benayoun et al. 2019). Furthermore, the activity of coagulation factors is associated with a higher risk of coronary heart disease (Lowe and Rumley 2014). However, we found no obvious histopathological lesions in the hearts or other organs (spleen, kidney, liver, lung) of non-breeders..

Therefore, if down-regulation of coagulation attentuates the aging process in *Fukomys*, it seems to exert its effect only, if at all, at a latter age not investigated here.

Two defense mechanisms were also found to be up-regulated in breeders: xenobiotic metabolism and TNF-α-signaling. Increased TNF-α-signaling often leads to the induction of apoptosis (Annibaldi and Meier 2018). In line with this, we found that apoptosis and P53-signaling were also up-regulated in breeders. Apoptosis is considered an important anti-cancer mechanism (Baig et al. 2016; Pistritto et al. 2016). We hypothesize this to compensate cancerogenic effects of the anabolic alterations described above (especially the up-regulation of the GH/IGF1 axis). In line with this, our more than 30 years’ breeding history with several *Fukomys* species in Germany and the Czech Republic suggests that breeders are as “cancer-proof” as non-breeders despite their much longer lifespan (own unpublished data and R. Šumbera, personal communication).

Many anabolic pathways are up-regulated in breeders across tissues: protein biosynthesis, myogenesis, and the GH/IGF1 axis. In line with this finding and with the fact that protein synthesis consumes 30 to 40% of a cell’s ATP budget (Hands et al. 2009), we observed increased expression of mitochondrial respiratory chain components, implying an increase in the capacity for cellular ATP production. On the other hand, protein degradation and clearance by the proteasome are also up-regulated in breeders. The fact that the expression of proteasomal genes is significantly linked to the genes of ribosome biogenesis and oxidative phosphorylation indicates that those processes influence, or even trigger, each other and hence are regulated in a coordinated manner in *Fukomys* mole-rats.

The results of differential expression of anabolic components such as the GH/IGF1 axis are surprising. They fall within a debate in aging research that has been highly controversial over time: based on the well-known fact that the expression and secretion of GH and IGF1 decline with age in humans and other mammals (Bartke 2019), Rudman *et al*. administered synthetic GH to elderly subjects in 1990, thereby reversing a number of aging-associated effects such as expansion of adipose mass (Rudman et al. 1990). This led to GH being celebrated as an anti-aging drug (Junnila et al. 2013), including dubious commercial offers. Today’s aging research, on the contrary, strongly assumes that the enhanced activity of the GH/IGF1 axis accelerates aging and that its suppression could extend lifespan even in humans (Lopez-Otin et al. 2013; Longo et al. 2015; Pitt and Kaeberlein 2015). In addition to several studies of synthetic GH in humans yielding less convincing results than those of Rudman *et al*., the main reasons for this turn are the results of studies on short-lived model organisms. From worms to mice, the impairment of the GH/IGF1 axis by genetic intervention consistently led to longer lifespans (Table 2), e.g., the up-regulation of Klotho - an IGF1 inhibitor - extended the mouse lifespan by as much as 30 % (Kurosu et al. 2005). As with the impairment of the GH/IGF1 axis, reducing of protein synthesis by decreasing the expression of MYC, a basal transcription factor, extended the mouse lifespan by as much as 20 % (Hofmann et al. 2015) whereas the impairment of the respiratory chain by rotenone resulted in prolongation of the killifish lifespan by 15 % (Baumgart et al. 2016) (Table 2).

**Table 2.** Behavior of important aging-relevant genes and pathways in this study.

| Gene/Pathway | Regulation in indicated direction and species can reduce lifespan | Regulation in indicated direction and species can extend lifespan | Differentially expressed in this study in indicated tissues and direction* | Gene/Pathway is expressed mainly in the following tissues |
| --- | --- | --- | --- | --- |
| Growth hormone (*GH1*) | Mouse ↑<sup>1</sup> | Mouse ↓<sup>2</sup>, Rat ↓<sup>2</sup> | Pituitary gland ↑ | Pituitary gland |
| Insulin growth factor 1 (*IGF1*) | – | Mouse ↓<sup>2</sup> | Adrenal gland ↑, Ovary ↑ | Liver, Ovary |
| IGF1-receptor (*IGF1R*) | – | Mouse ↓<sup>1</sup>, Worm ↓<sup>3</sup>, Fly ↓<sup>3</sup> | Adrenal gland ↓, Ovary ↓ | Many |
| Klotho (*KL*) | Mouse ↓<sup>3</sup> | Mouse ↑<sup>3</sup>, Worm ↑<sup>3</sup> | Ovary ↓ | Endocrine tissue, Kidney |
| *SIRT1* | – | Mouse ↑<sup>6</sup>, Worm ↑<sup>7</sup>, Fly ↑<sup>7</sup> | – | All |
| *MYC* | – | Mouse ↓<sup>8</sup> | Thyroid ↑ | All |
| *mTOR* | – | Mouse ↓<sup>9</sup>, Worm ↓<sup>9</sup>, Fly ↓<sup>9</sup> | Adrenal gland ↓ | All |
| *PRKAA2* (AMPK) | – | Worm ↑<sup>9</sup> | – | Many |
| *TP53* | Mouse ↑<sup>11</sup>, Fly ↑<sup>11</sup> | Worm ↓<sup>11</sup>, Mouse ↑<sup>11</sup>, Fly ↑<sup>11</sup> | – | All |
| *SOD2* | – | Worm ↓<sup>10</sup>, Fly ↑<sup>10</sup> | – | All |
| *FOXO3* | Fly ↑<sup>12</sup> | – | Ovary ↓, Adrenal gland ↓ | All |
| Protein synthesis | – | Mouse ↓<sup>8</sup>, Worm ↓<sup>9</sup>, Fly ↓<sup>9</sup> | Many ↑ | All |
| Proteasome | – | Worm ↑<sup>14</sup>, Fly ↑<sup>14</sup> | Gonads ↑, Adrenal gland ↑ | All |
| Lysosome | – | Worm ↑<sup>13</sup> | – | All |
| Respiratory chain | – | Worm ↓<sup>15</sup>, Fly ↓<sup>16</sup>, Killifish ↓<sup>17</sup> | Gonads ↑, Adrenal gland ↑, Blood ↑, Spleen ↑ | All |
| Apoptosis | Mouse ↑<sup>11</sup>, Fly ↑<sup>11</sup> | Worm ↓<sup>11</sup>, Mouse ↑<sup>11</sup>, Fly ↑<sup>11</sup> | Skin ↑, Pituitary gland ↑ | All |

1 (Junnila et al. 2013), 2 (Bartke 2019), 3 (van Heemst 2010), 4 (Elibol and Kilic 2018), 5 (Kwon et al. 2015), 6 (Sato et al. 2013), 7 (Burnett et al. 2011), 8 (Hofmann et al. 2015), 9 (Johnson et al. 2013), 10 (Edrey and Salmon 2014), 11 (Feng et al. 2011), 12 (Giannakou et al. 2004), 13 (Carmona-Gutierrez et al. 2016), 14 (Saez and Vilchez 2014), 15 (Dillin et al. 2002), 16 (Copeland et al. 2009), 17 (Baumgart et al. 2016)
\*Direction: breeder/non-breeder
\*\* Direction: old/young
– Either not affecting lifespan or not known to the best of our knowledge (column 1 and 2); no change (column 3)

Therefore, it is astonishing that massive up-regulation of these anabolic key components accompanies a lifespan extension of approximately 100 % in long-lived mammals and potentially even contributes to it. Several points could help to resolve this apparent contradiction: First, the up-regulation of anabolic pathways and key genes is at least partially accompanied by the regulation of other mechanisms that could plausibly compensate for deleterious effects. For example, it is, widely assumed that the negative impact of enhanced protein synthesis on lifespan is to a large extent caused by the accumulation of damaged or misfolded proteins, that is also known to contribute to aging-associated neurodegenerative diseases (Hipkiss 2007; Saez and Vilchez 2014; Carmona and Michan 2016). Up-regulation of the proteasome, as we observed in breeders in a weighted cross-tissue approach and especially in endocrine tissues, is known to counteract these effects by clearing damaged proteins leading to lifespan extension in worm and fly (Saez and Vilchez 2014). Enhanced proteasome activity has also been linked with higher longevity of the naked mole-rat compared to the laboratory mouse (Perez et al. 2009; Rodriguez et al. 2012) and in comparative approaches across several mammalian lineages (Pride et al. 2015). We hypothesize that the simultaneously high anabolic synthesis and catabolic degradation of proteins will lead to a higher protein turnover rate in breeders and, accompanied with that, a reduced accumulation of damaged and misfolded proteins. Similarly, it seems plausible that the up-regulation of the mitochondrial respiratory chain (oxidative phosphorylation) in breeders is compensated for by simultaneous up-regulation of the reactive oxygen hallmark: the mitochondrial respiratory chain is the main source of cellular ROS which can damage DNA, proteins and other cellular components (Balaban et al. 2005; Starkov 2008); the reactive oxygen hallmark consists by definition of genes that are known to be up-regulated in response to ROS treatments. Unsurprisingly, at least 25 % of these genes code for typical antioxidant enzymes such as thioredoxin, superoxide dismutase, peroxiredoxin, or catalase that can detoxify ROS. Furthermore, the known cancer promoting effects of an enhanced GH/IGF1 axis (Junnila et al. 2013) could, to some extent, be compensated for by up-regulation of apoptosis and p53 signaling, because these are major mechanisms of cancer suppression (Bieging et al. 2014). More generally, potential lifespan-extending effects of moderate up-regulation of both the GH/IGF1 axis and ROS production can also be viewed in the light of the hormesis hypothesis (Ristow 2014), which postulates that mild stressors can induce overall beneficial adaptive stress responses. In line with these arguments, we found higher resting metabolic rates in breeders compared to non-breeders in *Fukomys anselli* (Schielke et al. 2017), a species closely related to *F. micklemi*.

A second point that could help to resolve this apparent contradiction concerns the time of intervention. The transition from non-breeder to breeder takes place in adulthood, when by far the largest portion of the growth process has already been completed. In contrast, genetic interventions aimed at prolonging the lifespan by inhibiting the GH/IGFH1 axis (Table 2) usually affect the organisms throughout their entire lifespan, including infancy and youth. Therefore, it is still under debate whether the up-regulation of translation and anabolic processes by the GH/IGF1 axis independently enhances growth and aging or enhances aging only secondarily as a consequence of accelerated growth (Bartke 2017). Our results are an argument for the latter.

A third point is the question of the transferability of knowledge obtained in one species to other species. Most insights into current aging research originate from very short-lived model organisms (Table 2). It is clear, on the other hand, that the observed effects of lifespan-prolonging interventions listed in Table 2 are by far the smallest in the model organisms with the relatively longest lifespans: mice and rats.

Compared to most other mammals, however, even mice and rats are short-lived. Given their body weight and an often-used correlation between body weight and lifespan across mammals, their observed maximum lifespan of about four years corresponds to only 51 % (for mice) and 32 % (for rats) of the expected maximum lifespan. In contrast, humans can live as much as 463% as long as they would be expected based on their body weight. This disparity makes them, given this particular model, the most extreme known non-flying mammal species regarding maximum longevity residual (Tacutu et al. 2013). According to current data, *Fukomys* mole-rats reach values of ca. 200 % and thus can be regarded to be closer to humans than to mice or rats in this respect. It is currently unclear to what degree aging mechanisms and lifespan-affecting interventions that were discovered in short-lived model species apply to organisms that are far more long-lived, as in our experiment (e.g. (Parker et al. 2004; Keller and Jemielity 2006)). It seems reasonable to hypothesize that specific interventions that prolong the lifespan of long-lived organisms may have no major effect in short-lived species, and vice versa. For example, it could be that changes suitable to prolong the life of species with a low cancer-risk (e.g., African mole-rats, blind mole rats) (Buffenstein 2008; Gorbunova et al. 2012; Delaney et al. 2013) would have no or only a marginal effect in laboratory mice, whose primary cause of natural death is cancer (Seluanov et al. 2018). Therefore, it may also be possible that those gene expression patterns caused by our lifespan-extending intervention in *Fukomys* mole-rats but contradicting the current knowledge obtained from short-lived organisms may highlight differences in aging mechanisms between short-lived and long-lived species.

As a fourth perspective, one could interpret the down-regulation of the GH/IGF1 axis in non-breeders as a byproduct of the apparent up-regulation of the HPA axis in non-breeders, that may well be adaptive in itself. In the wild, non-breeding *Fukomys* mole-rats can maximize their inclusive fitness by either supporting their kin in their natal family or by founding a new family elsewhere. It has therefore been suggested that the shorter lives of non-breeders could be adaptive on the ultimate level if longevity were traded against some other fitness traits, such as competitiveness, to defend colonies against intruders or to enhance their chances for successful dispersal (Dammann and Burda 2006). A constitutively more activated HPA stress axis is expected to offer advantages for both family defense and dispersal but it carries the risk of status-specific aging symptoms, such as muscle weakness, lower GH/IGF1 axis activity, lower bone density etc. in the long run (Ferrau and Korbonits 2015). This effect may become even more pronounced under laboratory conditions where grown non-breeders cannot decide to disperse even if they would like to.

However, even the down-regulation of the GH/IGF-axis may be adaptive in itself for non-breeders if it has the potential to protect them from further damage. Note that for today’s conventional view that stronger activation of the GH/IGF1 axis accelerates aging (Table 2), it is generally challenging that decreasing activity is well documented to correlate with chronological age in a wide range of mammals, including mice, rats, dogs, and humans (Bartke 2019). Also, decreasing activity correlates, under pathologic conditions such as Cushing’s syndrome, with many symptoms akin to aging (Fig. 5). It has been suggested that one solution to this apparent contradiction may be that that the GH/IGF1 axis is adaptively down-regulated in aging organisms as a reaction to already accumulated aging-related symptoms so that additional damage can be avoided (Berryman et al. 2008; Milman et al. 2016).

Finally, recent findings of positively selected genes in African mole-rats (family Bathyergidae, containing also *Fukomys* and *Heterocephalus*) could partly explain some of the surprising results. It is striking that translation, and oxidative phosphorylation were among the strongest differentially expressed molecular processes concering the breeding status. Earlier, these processes were also reported to be the most affected by positive selection in the phylogeny of African mole-rats (Sahm et al. 2018b). Furthermore, *IGF1* was one of thirteen genes that were found to be under positive selection in the last common ancestor of the mole-rats. This may indicate that the corresponding mechanisms were evolutionarily adapted to be less detrimental and make their up-regulation more compatible with a long lifespan. Since the mere fact of positive selection does not permit to draw conclusions about the direction of the mechanistic effect, this hypothesis, however, needs to remain speculative.

## Conclusions

We performed a comprehensive transcriptome analysis that, for the first time within mammalian species, compared naturally occurring cohorts of species with massively diverging aging rates. The comparison of faster-aging *Fukomys* non-breeders with similar animals that were experimentally elevated to the slower-aging breeder status revealed by far the most robust transcriptome differences within endocrine tissue: adrenal gland, ovary, thyroid, and pituitary gland. Genes and pathways involved in anabolism, such as *GH*, *IGF1*, translation and oxidative phosphorylation, were differentially expressed. Their inhibition is among the best-documented life-prolonging interventions in a wide range of short-lived model organisms (Table 2). Surprisingly, however, we found that the expression of these mechanisms was consistently higher in slower-aging breeders than in faster-aging non-breeders. This indicates that even basic molecular mechanisms of the aging process known from short-lived species cannot easily be transferred to long-lived species. In particular, this applies to the role of the GH/IGF1 axis, which has in recent years been unilaterally described as harmful (Lopez-Otin et al. 2013; Pitt and Kaeberlein 2015; Bartke 2017). In addition, special features of the mole-rats could also contribute to the explanation of the unexpected result that genes and processes differentially expressed between reproductive statuses were also strongly altered during the evolution of the mole-rats (Sahm et al. 2018b). Another intriguing possibility is that, in line with the hormesis hypothesis (Ristow 2014), moderate harmful effects of anabolic processes can be hyper-compensated for by up-regulation of pathways such as proteasomes, P53-signaling and antioxidant defense against ROS that we observed in slower-aging breeders as well.

Furthermore, our work provides evidence that the HPA stress axis is a key regulator for the observed downstream effects, including the lifespan difference. The effects are likely to be triggered by differential expression of the gene *MC2R* coding for the ACTH receptor resulting in an altered stress response in breeders vs. non-breeders. This is supported by the fact that cortisol levels in the non-breeders are elevated. Furthermore, the set of direct and indirect target genes of the glucocorticoid receptors is strongly affected by differential expression, and numerous known downstream effects of glucocorticoid excess have been demonstrated for non-breeders, such as muscle weakness, weight gain, and GH/IGF1 axis impairment. Overall, this evidence suggests that *MC2R* and other genes along the described signalling pathway are promising targets for possible interventions in aging research.

## Methods

### Animal care and sampling

All animals were housed in glass terraria with dimensions adjusted to the size of the family (min. 40 cm × 60 cm) in the Department of General Zoology, Faculty of Biosciences, University of Duisburg-Essen. The terraria are filled with a 5 cm layer of horticultural peat or sawdust. Tissue paper strips, tubes, and solid shelters were provided as bedding/nesting materials and environmental enrichment. Potatoes and carrots are supplied ad libitum as stable food, supplemented with apples, lettuce, and cereals. *Fukomys* mole-rats do not drink free water. Temperature was kept fairly constant at 26± 1 °C, and humidity, at approximately 40 %. The daily rhythm was set to 12 hours darkness and 12 hours light.

New breeder pairs (new families) were established between March and May 2014. Each new family was founded by two unfamiliar, randomly chosen adult non-breeders of similar age (min/max/mean: 1.56/6.5/3.58 years in *F. mechowii*; 1.8/5.4/3.1 years in *F. micklemi*) and opposite sex and were taken from already existing separate colonies. These founder animals were moved to a new terrarium in which they were permanently mated. In both species, more than 80 % of these new pairs reproduced within the first 12 months (total number of offspring by the end of the year 2016, 82 *F. micklemi* and 81 *F. mechowii*). Only founders with offspring were subsequently assigned to the breeder cohort; founders without offspring were excluded from the study. Non-breeders remained in their natal family together with both parents and other siblings.

*F. mechowii* were sampled in five distinct sampling sessions between March 2015 and winter 2016/2017. F. *micklemi* were sampled in three distinct sampling sessions between November 2016 and July 2017. In both species, females were killed 4 to 6 months later than their male mates to ensure that these breeder females were neither pregnant nor lactating at the time of sampling, in order to exclude additional uncontrolled variables.

Before sampling, animals were anaesthetized with 6 mg/kg ketamine combined with 2.5 mg/kg xylazine (Garcia Montero et al. 2015). Once the animals lost their pedal withdrawal reflex, 1 to 2 ml of blood was collected by cardiac puncture, and the animals were killed by surgical decapitation. Blood samples (100 µl) were collected in RNAprotect Animal Blood reagent (Qiagen, Venlo, Netherlands). Tissue samples – hippocampus, hypothalamus, pituitary gland, thyroid, heart, skeletal muscle (M. quadriceps femoris), lateral skin, small intestine (ileum), upper colon, spleen, liver, kidney, adrenal gland, testis, and ovary – were transferred to RNAlater (Qiagen, Venlo, Netherlands) immediately after dissection and, following incubation, were stored at -80°C until analysis.

Animal housing and tissue collection were compliant with national and state legislation (breeding allowances 32-2-1180-71/328 and 32-2-11-80-71/345; ethics/animal experimentation approval 84– 02.04.2013/A164, Landesamt für Natur-, Umwelt- und Verbraucherschutz Nordrhein-Westfalen).

### RNA preparation and sequencing

For all tissues except blood, RNA was purified with the RNeasy Mini Kit (Qiagen) according to the manufacturer’s protocol. Blood RNA was purified with the RNeasy Protect Animal Blood Kit (Qiagen). Kidney and heart samples were treated with proteinase K before extraction, as recommended by the manufacturer. Library preparation was performed using the TruSeq RNA v2 kit (Illumina, San Diego, USA) which includes selection of poly-adenylated RNA molecules. RNA-seq was performed by single-end sequencing with 51 cycles in high-output mode on a HiSeq 2500 sequencing system (Illumina) and with at least 20 million reads per sample, as described in Table S9. Read data for *F. mechowii* and *F. micklemi* were deposited as European Nucleotide Archive study with the ID PRJEB29798 (Table S9).

### Read mapping and quantification

It was ensured for all samples that the results of the respective sequencing passed “per base” and “per sequence” quality checks of FASTQC (Andrews). The reads were then mapped against previously published and with human gene symbols annotated *F. mechowii* and *F. micklemi* transcriptome data (Sahm et al. 2018a; Sahm et al. 2018b). For both species, only the longest transcript isoform per gene was used; this is the method of choice for selecting a representative variant in large-scale experiments (Ezkurdia et al. 2015) (Data S2, S3). This selection resulted in 15,864 reference transcripts (genes) for *F. mechowii* and in 16,400 for *F. micklemi*. After mapping and quantification, we further analyzed only those reference transcripts whose gene symbols were present in the transcript catalogs of both species – this was the case for 15,199 transcripts (the size of the union was 17,065). As mapping algorithm “bwa aln” of the Burrows-Wheeler Aligner (BWA) (Li and Durbin 2009) was used, allowing no gaps and a maximum of two mismatches in the alignment. Only those reads that could be uniquely mapped to the respective gene were used for quantification. Read counts per gene and sample can be found in Data S4. As another check, we ensured that all samples exhibited a Pearson correlation coefficient of at least 90% in a pairwise comparison based on log2-transformed read counts against all other samples of the same experimental group as defined by samples that were equal in the tissue as well as the species, sex, and reproductive status of the source animal.

### Differentially expressed genes analysis

P-values for differential gene expression and fold-changes were determined with DESeq2 (Love et al. 2014) and a multi-factorial design. The DESeq2-algorithm also includes strict filtering based on a normalized mean gene count that makes further pre-filtering unnecessary (Love et al. 2014). Therefore, those genes whose read count was zero for all examined samples were removed before further analysis, thereby reducing the number of analyzed genes to 15,181. The multi-factorial design means that, separately for each tissue, we input the read count data of samples across species, sex, and reproductive status into DESeq2 for each sample. This allowed DESeq2 to perform DEG-analysis between the two possible states of each of the variables by controlling for additional variance in the other two variables. This approach resulted in a four-times higher sample size than with an approach that would have been based on comparisons of two experimental groups, each of which would be equal in tissue, species, sex, and reproductive status. It is known that the statistical power in RNA-seq experiments can increase considerably with sample size (Ching et al. 2014). P-values were corrected for multiple testing with the Benjamini-Hochberg correction (Benjamini and Y. 1995) (false discovery rate – FDR).

The results of the DEG-analysis can be found in Data S5-S7.

### Enrichment analysis on pathway and cross-tissue level

Let 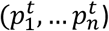 represent the p-values obtained from differential gene expression analysis in the tissue corresponding with index *t* and the indices 1, …, *n* corresponding to the examined genes. Furthermore, let 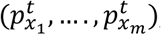, with 1 ≤ *x_i_* ≤ *n* and 1 ≤ *i* ≤ *m*, represent the p-values of genes with the indices *X* = *x*_1_, …, *x*_*m*_) belonging to a corresponding pathway that is tested for enrichment of differential expression signals. To determine the enrichment p-values at the pathway-level, we calculated the test statistic 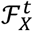 for the gene indices *X* in tissue index *t* according to Fisher’s method for combining p-values also known as Brown’s method:

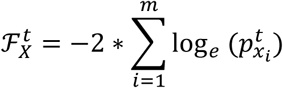

Because p-values at the gene level were, as is common in RNA-seq experiments (Väremo et al. 2013), not equally distributed, we empirically estimated the null distribution of the test statistic for each pathway instead of using the χ^2^ -distribution as frequently suggested in the literature (Fridley et al. 2010; Evangelou et al. 2012; Poole et al. 2016). This was done by calculating 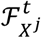 for 1,000 random drawings, each without replacement, 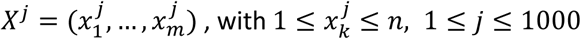. If the resulting p-value was zero (meaning 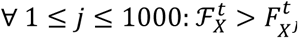), the procedure was repeated with 10,000 and 100,000 *X* were divided into *X_up_* and *X_down_* depending on whether their fold-change was > 1 or < 1 in breeder vs. non-breeder comparison, and 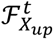 and 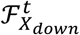 calculated. The ratio 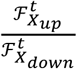 was used as an indicator for functional up- or down-regulation of the corresponding pathway (Fig. 4, Fig. S3, S4). Using this approach, enrichment p-values were estimated for all KEGG-pathways (Kanehisa et al. 2017) and MSigDB-hallmarks (Liberzon et al. 2015), as well as across all examined tissues (Data S8, S9). In addition, the procedure was applied to test whether the known 300 direct and 1300 indirect glucocorticoid receptor target genes (Phuc Le et al. 2005) were enriched for status-dependent differential expression signals (Tables S7, S8).

Similarly, cross-tissue DEG-p-values were weighted with a modified test statistic (c.f. (Heard and Rubin-Delanchy 2018)). Given the definitions from above, we calculated the weighted cross-tissue test statistic Ƒ^*g*^ for the gene *g* as follows:

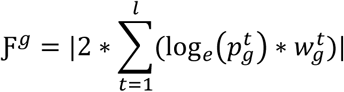

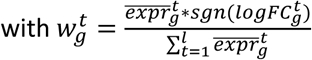

where 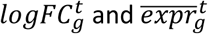 are the logarithmized fold-change between reproductive states and normalized mean expression (across sexes, species, and reproductive status) for the gene with index *g* and tissue with index *t* – both calculated by DESeq2 (Love et al. 2014) –, *sgn* is the signum function, and *l* is the number of examined tissues. The reasoning for using this test statistic was as follows: For the weighted cross-tissue analysis, we assumed that the gene serves the same function throughout the entire organism. Therefore, the test statistic given above weights the p-values of the various tissues by the respective expression levels in those tissues. This ensures that, weighting e. g., a ubiquitously expressed genes such as *TP53* is relatively equally across tissues, whereas for typical steroid hormone-biosynthesis genes, such as *CYP11A1*, the endocrine tissue results determine almost exclusively the weighted cross-tissue p-value. Furthermore, the test statistic rewards, based on the mentioned assumption, consistency in the direction of gene regulation throughout tissues. All calculated values for 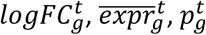, as well as the resulting Ƒ^*g*^ and p-values, can be found in Data S10.

Finally, weighted cross-tissue enrichment p-values at the pathway level were estimated by applying the above-described method at the pathway level (based on test statistic ℱ) to the gene level weighted cross-tissue p-values.

P-values were corrected for multiple testing with the Benjamini-Hochberg correction (Benjamini and Y. 1995) (FDR).

### Weighted gene co-expression network analysis

We used the WGCNA R package to perform weighted correlation network analysis (Langfelder and Horvath 2008) of all 636 samples at once. We followed the authors’ usage recommendation by choosing a soft power threshold based on scale-free topology and mean connectivity development (we chose power=26 with a soft R^2^ of 0.92 and a mean connectivity of 38.6), using biweight midcorrelation, setting maxPOutliers to 0.1, and using “signed” both as network and topological overlap matrix type. The maximum block size was chosen such that the analysis was performed with a single block and the minimum module size was set to 30. The analysis divided the genes into 26 modules, of which 5 were enriched for reproductive status DEGs based on Fisher’s exact test and an FDR threshold of 0.05. Those 5 modules were tested for enrichment among KEGG-pathways (Kanehisa et al. 2017) with the same test and significance threshold (Table S10). In addition, module eigengenes were determined and clustered (Table S10). Then the topological overlap matrix that resulted from the WGCNA analysis (*TOM* = [*tom*_*i,j*_], where the row indices 1 ≤ *i* ≤ |*examined genes*| correspond to genes and the column indices 1 ≤ *j* ≤ |*examined genes*| correspond to samples) was used to determine pairwise connectivity between all KEGG-pathways that showed differential expression at the weighted cross-tissue level (Fig. S5). Based on the definition of connectivity of genes in a WGCNA analysis (Langfelder and Horvath 2008), we defined the connectivity between two sets of indices *X* and *Y* each corresponding to genes as *k*_*X,Y*_ = ∑_*x* ∈ X\Y_ ∑_*x* ∈ Y\X_ *tom*_*x,y*_. P-values for the connectivities were determined against null distributions that were empirically estimated by determining for each pair *X* and *Y* the connectivities of 10,000 pairs of each |*X*| and |*Y*| randomly drawn indices (without replacement), respectively. Since “signed” was used as the network and topological overlap matrix type, the tests were one-sided.

### Other analysis steps

Hierarchical clustering (Fig. S2) was performed based on Pearson correlation coefficients of log2-transformed read counts between all sample pairs using the complete-linkage method (Defays 1977).

The principal variant component analysis (Fig. 2A) was performed with the pvca package from Bioconductor (Bushel 2013) and a minimum demanded percentile value of the amount of the variabilities, that the selected principal components needs to explain, of 0.5. Enrichments of DEGs among genes enlisted in the Digital Aging Atlas database ((Craig et al. 2015), Data S11) were determined with Fisher’s exact test, the Benjamini-Hochberg method (FDR) (Benjamini and Y. 1995) for multiple test correction, and a significance threshold of 0.05. Pathway visualization (Fig. S6-S9) was performed with Pathview (Luo and Brouwer 2013). For the direction analysis of *Fukomys* reproductive status DEGs in previous experiments in naked-mole rats and guinea pigs, we examined those 10 tissues that were examined in all species (Table S6, Data S12). Separately for each tissue and combination of species – naked mole-rat or guinea pig – and sex, we determined how many *Fukomys* reproductive status DEGs were up-regulated or down-regulated. We also performed two-sided binomial tests on each of these number pairs with a hypothesized success probability of 0.5. Furthermore, for each combination of species and sex, two-sided exact binomial tests using 0.5 as parameter were performed based on the sums of up-regulated and down-regulated genes across tissues (Table S6). For enrichment analysis of direct and indirect glucocorticoid receptor target genes, mouse mRNA RefSeq IDs from Phuc Le et al. (Phuc Le et al. 2005) were translated to human Entrez IDs and gene symbols via Ensembl Biomart (Data S13). To statistically analyze the weight gain of the animals during the experiment, we used a type II ANOVA with status, species, sex, and age as independent variables (Table S2); the weight gain, defined as the difference in weights at beginning and the end of the experiment, as dependent variable (Fig. 6B); and no interaction terms. If interaction terms were also used for the model, the p-value for the difference in means between breeders and non-breeders changed from 7.49*10^-3^, as reported above, to 7.46*10^-6^.

### Bone density measurements

Frozen carcasses of all *F. micklemi* that had been part of the transcriptome study were scanned with a self-shielded desktop small-animal computer tomography scanner (X-CUBE, Molecubes, Belgium). The x-ray source was a tungsten anode (peak voltage, 50 kVp; tube current, 350 µA; 0.8 mm aluminum filter). The detector was a cesium iodide (CsI) flat-panel, building up a screen with 1536 x 864 pixels. Measurements were carried for individual 120 ms exposures, with angular sampling intervals of 940 exposures per rotation, for to a total of 7 rotations and a total exposure time of 789.6 seconds.

First, we first performed a calibration of the reconstructed CT data in terms of equivalent mineral density. For this purpose, we used a bone density calibration phantom (BDC; QRM GmbH, Moehrendorf, Germany) composed of five cylindrical inserts with a diameter of 5 mm containing various densities of calcium hydroxyapatite (CaHA) surrounded by epoxy resin on a cylindrical shape. The nominal values of CaHA were 0, 100, 200, 400, and 800 mg HA/cm^3^, corresponding to a density of 1.13, 1.16, 1.25, 1.64, and 1.90 g/cm^3^ (certified with an accuracy of ± 0.5 %). The BDC was imaged and reconstructed with the same specifications as each probe. From the reconstructed Hounsfield Units, a linear relationship was determined against the known mineral concentrations.

Reconstruction of the acquired computer tomography data was carried with an Image Space Reconstruction Algorithm, and spatial resolution was limited to the 100 µm voxel matrix reconstruction. Spherical regions of interest (radius, 0.7 mm) were drawn on the sagittal plane of vertebrae T12, L1, and L2. Care was taken to include all cancellous bone, excluding the cortical edges. Average Hounsfield Unit values were computed on the calibration curve to finally retrieve equivalent densities of the regions of interest.

Statistical analysis was performed using general linear models with bone density (Hounsfield Units) as dependent variable, age (in days) as continuous covariate, and reproductive status and sex as nominal cofactors. Models were calculated for each vertebra individually (individual models) and across all three vertebrae (full model); in this latter case, vertebral number was added as additional categorial cofactor.

In all models, only main effects were calculated, no interactions. Analyses were performed with IBM SPSS version 25 (Fig. 6C, Table S11).

## Abbreviations

ANOV: Aanalysis of variance
ACTH: adrenocorticotropic hormone
DAA: Digital Aging Atlas
DEG: differentially expressed gene
DHEA: Dehydroepiandrosterone
FDR: false discovery rate
KEGG: Kyoto Encyclopedia of Genes and Genomes
MSigDB: Molecular Signatures Database

## Data availability

Read datasets generated during the current study are available in the European Nucleotide Archive, study ID: PRJEB29798.

## Acknowledgments

We thank Ivonne Görlich, Christiane Vole, and Klaus Huse for excellent assistance in the preparation of biological samples. We thank Konstantin Riege for fruitful discussions on the analysis of the data. We thank Debra Weih and Flo Witte for proofreading the manuscript.

## Authors’ contributions

PD, MP, and KS conceived the project (with input from SB) and acquired the funding. PD, KS, and AS monitored the implementation of the project. PD, SB, HB, and PVD were responsible for housing the animals and implementing concrete mating schemes. PD, SB, ST, MGo, MS, JK, and PFC performed physiological measurements. PD, YH, KS, AS, SB, and MB sampled the animals. MG supervised library preparation and sequencing. AS, KS, and MB analyzed the data. AS, PD, MP, KS, SH, and PK interpreted the data and drafted the manuscript concept. All authors wrote and approved the manuscript.

## Ethics declarations

The authors declare no competing interests. Animal housing and tissue collection were compliant with national and state legislation (breeding allowances 32-2-1180-71/328 and 32-2-11-80-71/345, ethics/animal experimentation approval 84– 02.04.2013/A164, Landesamt für Natur-, Umwelt- und Verbraucherschutz Nordrhein-Westfalen).

## Supplementary information

### Supplement Figures

**Figure S1.** Overview of the tissues examined in this study with a schematic mole-rat representation.

**Figure S2.** Clustering of the 636 examined samples. Clustering was performed based on the Euclidian distance of logarithmized pairwise Pearson read count correlations using the UPGMA method.

**Figure S3.** Full overview of Kyoto Encyclopedia of Genes and Genomes (KEGG) pathways that are enriched for status-dependent differential gene expression.

**Figure S4.** Full overview of Molecular Signatures Database (MsigDB) hallmarks that are enriched for status-dependent differential gene expression.

**Figure S5.** Pairwise connectivity of metabolic pathways that are enriched for status-dependent differential gene expression at the cross-tissue level.

**Figure S6.** Status-dependent regulation of Kyoto Encyclopedia of Genes and Genomes (KEGG) pathway hsa00140 Steroid hormone biosynthesis at the cross-tissue level.

**Figure S7.** Status-dependent regulation of Kyoto Encyclopedia of Genes and Genomes (KEGG) pathway hsa05016 Huntington’s disease at the cross-tissue level.

**Figure S8.** Status-dependent regulation of Kyoto Encyclopedia of Genes and Genomes (KEGG) pathway hsa05012 Parkinson’s disease at the cross-tissue level.

**Figure S9.** Status-dependent regulation of Kyoto Encyclopedia of Genes and Genomes (KEGG) pathway hsa050160 Alzheimer’s disease at the cross-tissue level.

### Supplement Tables

**Table S1.** Overview of number of samples that were examined with regard to status, sex, species, and tissue.

**Table S2.** Animal description.

**Table S3.** Overview of the tissues that were successfully sampled for each type of animal.

**Table S4.** Pairing scheme: *F. mechowii*.

**Table S5.** Pairing scheme: *F. micklemi*.

**Table S6.** Analysis of the direction of status-dependent differentially expressed genes that were identified in this study (two *Fukomys* species) in similar experiments with naked mole-rats and guinea pigs (Bens et al. 2018, https://www.ncbi.nlm.nih.gov/pmc/articles/PMC6090939/).

**Table S7.** Analysis of status-dependent differential gene expression enrichment on glucocorticoid receptor target genes that were determined by Phuc Le at al. 2005 (https://www.ncbi.nlm.nih.gov/pmc/articles/PMC1186734/), using chromatin immunoprecipitation (CHIP).

**Table S8.** Analysis of status-dependent differential gene expression enrichment on glucocorticoid receptor target genes that were determined by Phuc Le at al. 2005 (https://www.ncbi.nlm.nih.gov/pmc/articles/PMC1186734/), using a differential gene expression analysis.

**Table S9.** Sample description.

**Table S10.** WGCNA module clustering and functional enrichment analysis regarding these modules.

**Table S11.** Bone density measurements.

### Supplement Data

**Data S1.** (zip) For each KEGG pathway and MSigDB hallmark that was detected to be significantly (FDR < 0.1) enriched for status-dependent differential expression at the weighted cross-tissue level, the data set contains a *.tsv file with the genes that form the respective pathway/hallmark sorted by their individual contribution to the enrichment. The files also provide an overview of the p-values and fold-changes of those genes in those tissues in which the genes are expressed most highly.

**Data S2.** (fa.gz) Transcript isoforms of *F. mechowii* used for read mapping.

**Data S3.** (fa.gz) Transcript isoforms of *F. micklemi* used for read mapping.

**Data S4.** (tsv.gz) Raw read counts for all 17,065 genes and 636 samples that were analyzed in this study using RNA-seq.

**Data S5.** (zip) DESeq2 results for status-dependent gene expression (direction: breeder/non-breeder). The data set contains one *.tsv file per analyzed tissue.

**Data S6.** (zip) DESeq2 results for sex-dependent gene expression (direction: female/male). The data set contains one *.tsv file per analyzed tissue.

**Data S7.** (zip) DESeq2 results for species-dependent gene expression (direction: mechowii/micklemi). The data set contains one *.tsv file per analyzed tissue.

**Data S8.** (zip) Enrichment analysis results for status-dependent gene expression on KEGG pathways. The data set contains one *.tsv file per analyzed tissue, as well as an additional *.tsv file for the weighted cross-tissue level results.

**Data S9.** (zip) Enrichment analysis results for status-dependent gene expression on MSigDB hallmarks. The data set contains one *.tsv file per analyzed tissue, as well as an additional *.tsv file for the weighted cross-tissue level results.

**Data S10.** (tsv.gz) Overview of the weighted cross-tissue differential gene expression analysis. Contains all p-values, fold-changes, and mean expression values for all genes across all tissues as well as the weighted, combined cross-tissue test statistics, p-values, and fold-changes for all genes.

**Data S11.** (tsv) Genes of the Digital Aging Atlas used in this study.

**Data S12.** (zip) Comparison of status-dependent differentially expressed genes with results of an earlier, similar study involving naked mole-rats (NMRs) and guinea pigs (GPs). The data set contains one *.tsv file for each of the ten tissues that were examined in both studies. Each *.tsv file lists the differentially expressed genes for the respective tissue as identified in this study with the determined FDRs and fold-changes, as well as the fold-changes determined in the earlier study using naked mole-rats and guinea pigs.

**Data S13**. (zip) Glucocorticoid receptor target genes that were determined by Phuc Le *et al*. and tested in this study for enrichment of status-dependent differential gene expression. The data set contains one *.tsv file each for target genes determined via chromatin immunoprecipitation (CHIP) and differential gene expression analysis. Phuc Le et al. determined differentially expressed genes between mice that were treated with exogenous glucocorticoids and untreated controls. The relevant table columns from Phuc Le et al. were added by human gene symbol and Entrez IDs that were used for enrichment analysis.

