## Supplement Figures S1-S9 for "The HPA stress axis shapes aging rates in long-lived, social mole-rats"

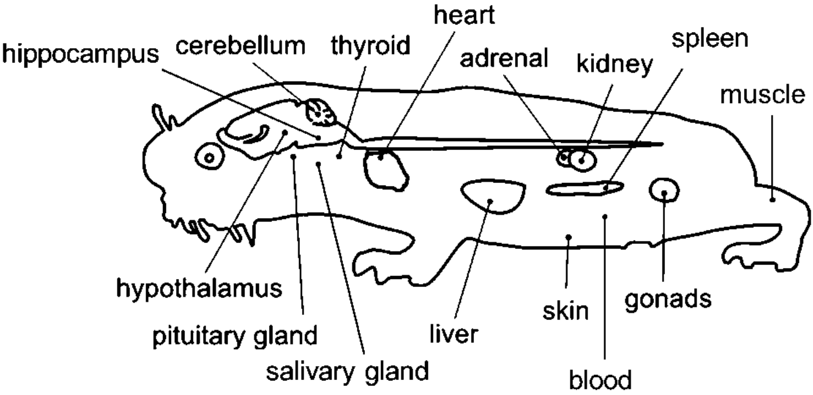


**Figure S1.** Overview of the tissues examined in this study with a schematic mole-rat representation.

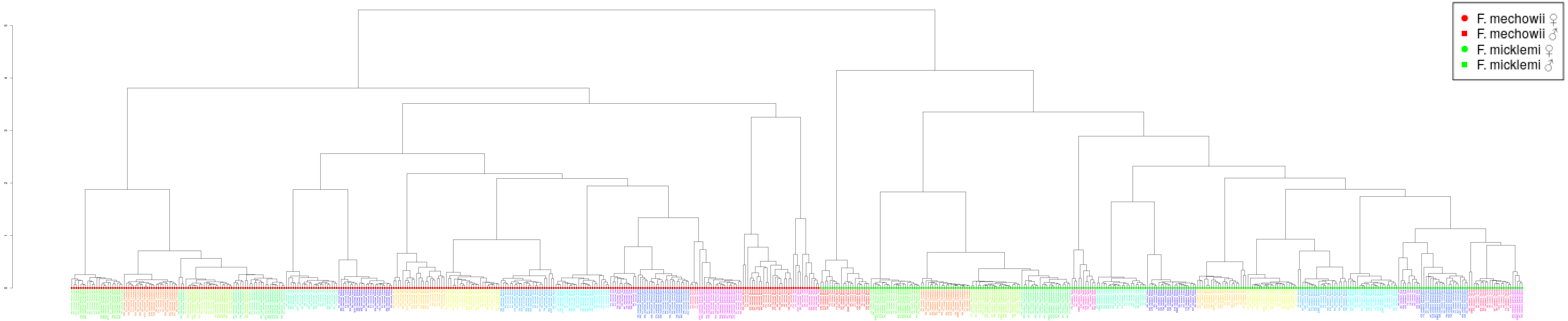


**Figure S2.** Clustering of the 636 examined samples. Clustering was performed based on the Euclidian distance of logarithmized pairwise Pearson read count correlations according to the unweighted pair group method with arithmetic mean (UPGMA). Bigger sample labels, breeders; smaller sample, non-breeders.

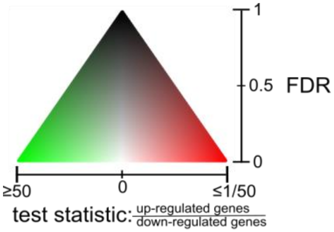


| Pathway | Cross−tissue | Adrenal gland | Blood | Cerebellum | Heart | Hippocampus | Hypothalamus | Kidney | Liver | Muscle | Ovar | Pituitary gland | Salivary gland | Skin | Spleen | Testis | Thyroid |
|---|---|---|---|---|---|---|---|---|---|---|---|---|---|---|---|---|---|
| hsa03010 Ribosome | 0.000 | 0.000 | 0.545 | 0.902 | 0.000 | 0.000 | 1.000 | 0.000 | 0.177 | 0.053 | 0.000 | 0.207 | 1.000 | 1.000 | 0.001 | 0.000 | 0.034 |
| hsa05016 Huntington's disease | 0.004 | 0.000 | 0.213 | 0.895 | 0.869 | 1.000 | 1.000 | 0.479 | 0.664 | 0.813 | 0.007 | 0.996 | 1.000 | 0.987 | 0.101 | 0.000 | 0.171 |
| hsa03050 Proteasome | 0.004 | 0.000 | 0.593 | 0.857 | 1.000 | 1.000 | 0.815 | 0.993 | 0.981 | 0.155 | 0.003 | 0.996 | 1.000 | 0.063 | 0.264 | 0.000 | 1.000 |
| hsa00140 Steroid hormone biosynthesis | 0.006 | 1.000 | 0.053 | 0.850 | 0.869 | 1.000 | 0.554 | 0.266 | 0.764 | 0.009 | 0.542 | 0.983 | 0.228 | 0.216 | 0.008 | 1.000 | 1.000 |
| hsa05012 Parkinson's disease | 0.018 | 0.000 | 0.053 | 0.883 | 1.000 | 0.684 | 1.000 | 0.098 | 0.991 | 0.868 | 0.002 | 0.996 | 1.000 | 0.929 | 0.118 | 0.002 | 0.263 |
| hsa03008 Ribosome biogenesis in eukaryotes | 0.023 | 0.094 | 0.655 | 0.814 | 0.725 | 1.000 | 0.652 | 0.975 | 0.380 | 0.524 | 0.722 | 0.996 | 0.982 | 0.987 | 0.118 | 0.713 | 0.388 |
| hsa04260 Cardiac muscle contraction | 0.031 | 0.408 | 0.602 | 0.850 | 0.598 | 1.000 | 0.968 | 0.820 | 0.931 | 0.367 | 0.605 | 0.163 | 0.982 | 0.932 | 0.149 | 0.537 | 0.538 |
| hsa05010 Alzheimer's disease | 0.031 | 0.000 | 0.087 | 0.857 | 0.889 | 1.000 | 0.919 | 0.076 | 0.897 | 0.650 | 0.019 | 0.996 | 0.982 | 0.889 | 0.239 | 0.009 | 0.263 |
| hsa00190 Oxidative phosphorylation | 0.033 | 0.000 | 0.593 | 0.914 | 0.954 | 1.000 | 1.000 | 0.813 | 0.991 | 0.402 | 0.002 | 0.996 | 1.000 | 0.682 | 0.239 | 0.011 | 0.263 |
| hsa04976 Bile secretion | 0.038 | 1.000 | 0.053 | 0.857 | 0.598 | 1.000 | 0.228 | 0.046 | 0.556 | 0.588 | 0.404 | 0.254 | 1.000 | 0.933 | 0.001 | 1.000 | 1.000 |
| hsa00982 Drug metabolism − cytochrome P450 | 0.038 | 1.000 | 0.046 | 0.857 | 0.869 | 1.000 | 0.736 | 0.998 | 0.057 | 0.301 | 0.966 | 0.996 | 0.982 | 0.553 | 0.000 | 0.824 | 1.000 |
| hsa04975 Fat digestion and absorption | 0.038 | 1.000 | 0.213 | 0.643 | 0.925 | 1.000 | 0.324 | 0.975 | 0.897 | 0.189 | 0.179 | 0.358 | 0.982 | 0.520 | 0.062 | 1.000 | 1.000 |
| hsa03013 RNA transport | 0.053 | 0.023 | 0.644 | 0.814 | 0.598 | 1.000 | 1.000 | 0.820 | 0.897 | 0.418 | 0.719 | 0.996 | 1.000 | 0.936 | 0.239 | 0.537 | 0.309 |
| hsa02010 ABC transporters | 0.081 | 1.000 | 0.644 | 0.046 | 0.725 | 1.000 | 0.794 | 0.205 | 0.687 | 0.406 | 0.687 | 0.923 | 1.000 | 0.429 | 0.662 | 0.924 | 0.370 |
| hsa04977 Vitamin digestion and absorption | 0.122 | 1.000 | 0.465 | 0.874 | 0.598 | 1.000 | 0.607 | 0.811 | 0.717 | 0.000 | 0.019 | 0.996 | 0.987 | 0.470 | 0.065 | 1.000 | 0.896 |
| hsa03060 Protein export | 0.174 | 0.094 | 0.919 | 0.814 | 0.171 | 1.000 | 0.968 | 0.000 | 0.981 | 0.545 | 0.752 | 0.475 | 0.982 | 0.400 | 0.587 | 0.391 | 1.000 |
| hsa00980 Metabolism of xenobiotics by cytochrome P450 | 0.177 | 1.000 | 0.160 | 0.927 | 0.869 | 1.000 | 0.919 | 0.998 | 0.122 | 0.100 | 0.829 | 0.996 | 0.982 | 0.179 | 0.023 | 0.924 | 1.000 |
| hsa01100 Metabolic pathways | 0.180 | 1.000 | 0.000 | 0.874 | 0.869 | 1.000 | 0.324 | 0.994 | 0.046 | 0.001 | 0.000 | 0.288 | 0.982 | 0.542 | 0.020 | 0.924 | 0.878 |
| hsa03320 PPAR signaling pathway | 0.208 | 1.000 | 0.010 | 0.860 | 0.725 | 1.000 | 0.481 | 0.912 | 0.046 | 0.100 | 1.000 | 0.912 | 0.982 | 0.932 | 0.000 | 1.000 | 0.263 |
| hsa00120 Primary bile acid biosynthesis | 0.208 | 1.000 | 0.046 | 0.702 | 0.869 | 1.000 | 0.228 | 0.959 | 0.690 | 0.169 | 0.858 | 0.912 | 0.982 | 0.453 | 0.010 | 1.000 | 1.000 |
| hsa00310 Lysine degradation | 0.270 | 0.076 | 0.669 | 0.857 | 0.860 | 1.000 | 0.338 | 0.998 | 0.616 | 0.253 | 0.719 | 0.588 | 0.982 | 0.961 | 0.568 | 0.033 | 1.000 |
| hsa00900 Terpenoid backbone biosynthesis | 0.270 | 0.772 | 0.521 | 0.858 | 0.869 | 1.000 | 0.827 | 0.975 | 0.839 | 0.301 | 0.035 | 0.856 | 1.000 | 0.939 | 0.559 | 0.924 | 1.000 |
| hsa03040 Spliceosome | 0.322 | 0.094 | 0.644 | 0.857 | 0.598 | 1.000 | 0.825 | 0.993 | 0.659 | 0.781 | 1.000 | 0.923 | 1.000 | 0.429 | 0.816 | 0.342 | 0.023 |
| hsa04610 Complement and coagulation cascades | 0.322 | 1.000 | 0.046 | 0.702 | 0.889 | 1.000 | 0.228 | 0.542 | 0.764 | 0.939 | 1.000 | 0.912 | 0.228 | 0.554 | 0.000 | 1.000 | 1.000 |
| hsa00330 Arginine and proline metabolism | 0.368 | 1.000 | 0.067 | 0.914 | 0.598 | 1.000 | 0.130 | 0.304 | 0.209 | 0.379 | 0.274 | 0.046 | 0.982 | 0.437 | 0.161 | 1.000 | 1.000 |
| hsa00030 Pentose phosphate pathway | 0.420 | 1.000 | 0.279 | 0.814 | 0.725 | 1.000 | 1.000 | 0.998 | 0.714 | 0.185 | 0.019 | 0.475 | 0.893 | 0.711 | 0.336 | 1.000 | 1.000 |
| hsa04146 Peroxisome | 0.561 | 1.000 | 0.279 | 0.877 | 0.598 | 1.000 | 0.740 | 0.998 | 0.363 | 0.011 | 0.612 | 0.996 | 0.982 | 0.992 | 0.662 | 0.830 | 0.896 |
| hsa05140 Leishmaniasis | 0.694 | 1.000 | 0.644 | 0.883 | 1.000 | 1.000 | 0.228 | 0.998 | 0.380 | 0.876 | 1.000 | 0.996 | 0.982 | 0.003 | 0.847 | 1.000 | 1.000 |
| hsa00360 Phenylalanine metabolism | 0.701 | 1.000 | 0.087 | 0.857 | 1.000 | 1.000 | 0.310 | 0.850 | 0.897 | 0.041 | 0.404 | 0.494 | 1.000 | 0.110 | 0.101 | 1.000 | 1.000 |
| hsa00380 Tryptophan metabolism | 0.704 | 1.000 | 0.053 | 0.874 | 0.869 | 1.000 | 0.374 | 0.998 | 0.046 | 0.205 | 1.000 | 0.475 | 0.947 | 0.484 | 0.366 | 1.000 | 1.000 |
| hsa00280 Valine, leucine and isoleucine degradation | 0.805 | 1.000 | 0.137 | 0.874 | 0.869 | 1.000 | 0.239 | 0.998 | 0.037 | 0.155 | 0.299 | 0.669 | 0.982 | 0.131 | 0.523 | 0.924 | 1.000 |
| hsa04380 Osteoclast differentiation | 0.805 | 1.000 | 0.646 | 0.997 | 1.000 | 1.000 | 0.470 | 0.998 | 0.690 | 0.620 | 1.000 | 0.912 | 0.982 | 0.002 | 0.553 | 0.996 | 1.000 |
| hsa00260 Glycine, serine and threonine metabolism | 0.813 | 1.000 | 0.053 | 0.857 | 0.920 | 1.000 | 0.228 | 0.850 | 0.104 | 0.041 | 0.983 | 0.465 | 0.982 | 0.531 | 0.076 | 1.000 | 1.000 |
| hsa01040 Biosynthesis of unsaturated fatty acids | 0.842 | 1.000 | 0.053 | 0.456 | 0.342 | 1.000 | 0.959 | 0.959 | 0.363 | 0.009 | 0.809 | 0.833 | 1.000 | 0.965 | 0.216 | 0.924 | 1.000 |
| hsa00071 Fatty acid metabolism | 0.842 | 1.000 | 0.277 | 0.944 | 0.598 | 1.000 | 0.200 | 0.998 | 0.209 | 0.009 | 0.444 | 0.494 | 0.982 | 0.933 | 0.568 | 0.824 | 0.651 |
| hsa00830 Retinol metabolism | 0.842 | 1.000 | 0.212 | 0.509 | 0.869 | 1.000 | 1.000 | 0.998 | 0.380 | 0.756 | 0.863 | 0.996 | 1.000 | 0.661 | 0.023 | 0.909 | 0.878 |
| hsa04960 Aldosterone−regulated sodium reabsorption | 0.842 | 0.967 | 0.330 | 0.914 | 0.869 | 1.000 | 0.541 | 0.795 | 0.997 | 0.189 | 0.004 | 0.813 | 0.982 | 0.798 | 0.152 | 0.947 | 0.651 |
| hsa00040 Pentose and glucuronate interconversions | 0.842 | 1.000 | 0.076 | 0.874 | 1.000 | 1.000 | 0.525 | 0.994 | 0.209 | 0.028 | 0.961 | 0.996 | 0.982 | 0.804 | 0.086 | 0.649 | 0.948 |
| hsa04910 Insulin signaling pathway | 0.897 | 0.266 | 0.602 | 0.914 | 0.598 | 1.000 | 0.415 | 0.912 | 0.122 | 0.397 | 0.856 | 0.447 | 1.000 | 0.987 | 0.577 | 0.537 | 0.046 |
| hsa04145 Phagosome | 0.919 | 1.000 | 0.605 | 0.513 | 1.000 | 1.000 | 0.255 | 0.795 | 0.999 | 0.948 | 0.444 | 0.967 | 0.982 | 0.033 | 0.816 | 0.929 | 1.000 |
| hsa05320 Autoimmune thyroid disease | 0.919 | 1.000 | 0.644 | 0.814 | 1.000 | 1.000 | 0.130 | 0.998 | 1.000 | 0.988 | 1.000 | 0.996 | 0.982 | 0.028 | 0.964 | 1.000 | 1.000 |
| hsa04974 Protein digestion and absorption | 0.965 | 1.000 | 0.465 | 0.944 | 1.000 | 1.000 | 0.607 | 0.542 | 0.972 | 0.190 | 0.019 | 0.254 | 0.982 | 0.932 | 0.152 | 1.000 | 1.000 |
| hsa04510 Focal adhesion | 0.972 | 1.000 | 0.605 | 0.771 | 0.598 | 1.000 | 0.130 | 0.046 | 0.981 | 0.556 | 0.722 | 0.200 | 1.000 | 0.484 | 0.559 | 0.909 | 1.000 |
| hsa00620 Pyruvate metabolism | 0.972 | 1.000 | 0.228 | 0.857 | 0.693 | 1.000 | 0.655 | 0.975 | 0.037 | 0.588 | 0.966 | 0.475 | 0.982 | 0.467 | 0.568 | 0.603 | 1.000 |
| hsa04512 ECM−receptor interaction | 0.972 | 1.000 | 0.093 | 0.902 | 0.725 | 1.000 | 0.228 | 0.005 | 0.999 | 0.726 | 0.809 | 0.632 | 0.987 | 0.553 | 0.559 | 0.947 | 1.000 |
| hsa00640 Propanoate metabolism | 0.972 | 1.000 | 0.271 | 0.877 | 0.828 | 1.000 | 0.481 | 0.975 | 0.037 | 0.402 | 0.745 | 0.682 | 0.982 | 0.391 | 0.865 | 0.924 | 1.000 |
| hsa04141 Protein processing in endoplasmic reticulum | 0.972 | 0.051 | 0.126 | 0.114 | 0.869 | 1.000 | 0.862 | 0.959 | 0.897 | 0.522 | 0.277 | 0.588 | 0.228 | 0.003 | 0.812 | 1.000 | 1.000 |
| hsa04010 MAPK signaling pathway | 0.972 | 1.000 | 0.324 | 0.643 | 0.920 | 1.000 | 0.607 | 0.912 | 0.695 | 0.305 | 1.000 | 0.254 | 1.000 | 0.021 | 0.296 | 0.824 | 1.000 |
| hsa00920 Sulfur metabolism | 0.972 | 1.000 | 0.639 | 0.874 | 0.998 | 1.000 | 1.000 | 0.975 | 0.823 | 0.020 | 0.843 | 0.799 | 0.982 | 0.798 | 0.264 | 0.909 | 0.896 |
| hsa05110 Vibrio cholerae infection | 0.972 | 0.182 | 0.988 | 0.857 | 1.000 | 1.000 | 0.900 | 0.850 | 0.839 | 0.211 | 0.009 | 0.848 | 0.982 | 0.553 | 0.559 | 1.000 | 1.000 |
| hsa04514 Cell adhesion molecules (CAMs) | 0.998 | 1.000 | 0.354 | 0.857 | 0.869 | 1.000 | 0.200 | 0.820 | 0.999 | 0.028 | 0.961 | 0.994 | 1.000 | 0.011 | 0.865 | 1.000 | 1.000 |
| hsa00410 beta−Alanine metabolism | 0.998 | 1.000 | 0.412 | 0.997 | 0.954 | 1.000 | 0.415 | 0.998 | 0.046 | 0.249 | 0.789 | 0.912 | 0.982 | 0.929 | 0.865 | 1.000 | 1.000 |
| hsa05323 Rheumatoid arthritis | 0.998 | 1.000 | 0.868 | 0.914 | 1.000 | 1.000 | 0.575 | 0.998 | 0.897 | 0.646 | 1.000 | 0.833 | 0.982 | 0.002 | 0.762 | 1.000 | 1.000 |
| hsa05160 Hepatitis C | 0.998 | 1.000 | 0.831 | 0.858 | 1.000 | 1.000 | 0.523 | 0.912 | 0.870 | 0.296 | 0.799 | 0.996 | 0.982 | 0.033 | 0.847 | 1.000 | 0.878 |
| hsa05142 Chagas disease (American trypanosomiasis) | 0.998 | 1.000 | 0.228 | 0.974 | 1.000 | 1.000 | 0.536 | 0.820 | 0.717 | 0.703 | 1.000 | 0.857 | 0.982 | 0.046 | 0.762 | 1.000 | 1.000 |
| hsa04612 Antigen processing and presentation | 0.998 | 1.000 | 0.599 | 0.645 | 1.000 | 1.000 | 0.327 | 1.000 | 1.000 | 0.650 | 1.000 | 0.996 | 0.982 | 0.002 | 1.000 | 0.947 | 1.000 |
| hsa05145 Toxoplasmosis | 0.998 | 1.000 | 0.723 | 0.857 | 1.000 | 1.000 | 0.228 | 0.998 | 0.841 | 0.878 | 1.000 | 0.843 | 0.982 | 0.021 | 0.981 | 1.000 | 1.000 |
| hsa04672 Intestinal immune network for IgA production | 0.998 | 1.000 | 0.912 | 0.857 | 1.000 | 1.000 | 0.481 | 0.998 | 0.999 | 0.769 | 1.000 | 0.996 | 0.982 | 0.003 | 0.947 | 1.000 | 1.000 |
| hsa04060 Cytokine−cytokine receptor interaction | 0.998 | 1.000 | 0.310 | 0.846 | 1.000 | 1.000 | 0.811 | 0.998 | 0.997 | 0.957 | 1.000 | 0.996 | 0.982 | 0.003 | 0.880 | 1.000 | 1.000 |
| hsa04660 T cell receptor signaling pathway | 0.998 | 1.000 | 0.639 | 0.979 | 1.000 | 1.000 | 0.737 | 0.998 | 0.976 | 0.650 | 0.918 | 0.996 | 1.000 | 0.021 | 0.745 | 0.924 | 0.948 |

**Figure S3.** Full overview of Kyoto Encyclopedia of Genes and Genomes (KEGG) pathways that are enriched for caste-dependent differential gene expression. Listed are all pathways that show a significant differential expression signal (FDR < 0.05) in at least one of the tissues or on the cross tissue-level (FDR < 0.1).

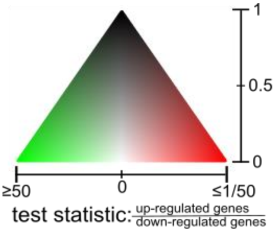


| | Cross–tissue | Adrenal gland | Blood | Cerebellum | Heart | Hippocampus | Hypothalamus | Kidney | Liver | Muscle | Ovar | Pituitary gland | Salivary gland | Skin | Spleen | Testis | Thyroid |
|---|---|---|---|---|---|---|---|---|---|---|---|---|---|---|---|---|---|
| HALLMARK_MYC_TARGETS_V1 | 0.000 | 0.005 | 0.027 | 0.998 | 0.011 | 0.863 | 0.989 | 0.165 | 0.030 | 0.870 | 0.030 | 0.843 | 0.936 | 0.476 | 0.007 | 0.060 | 0.617 |
| HALLMARK_XENOBIOTIC_METABOLISM | 0.000 | 1.000 | 0.000 | 0.998 | 0.225 | 0.863 | 0.362 | 0.165 | 0.050 | 0.025 | 0.050 | 0.468 | 0.409 | 0.039 | 0.000 | 1.000 | 0.998 |
| HALLMARK_OXIDATIVE_PHOSPHORYLATION | 0.008 | 0.000 | 0.025 | 0.998 | 0.233 | 0.379 | 0.989 | 0.213 | 0.331 | 0.153 | 0.000 | 1.000 | 0.901 | 0.980 | 0.007 | 0.003 | 0.617 |
| HALLMARK_HYPOXIA | 0.012 | 0.896 | 0.014 | 0.998 | 0.225 | 0.423 | 0.989 | 0.956 | 0.135 | 0.784 | 0.050 | 0.057 | 0.550 | 0.000 | 0.153 | 1.000 | 0.780 |
| HALLMARK_COAGULATION | 0.020 | 1.000 | 0.000 | 0.998 | 0.450 | 0.921 | 0.937 | 0.274 | 0.094 | 0.308 | 0.888 | 0.820 | 0.969 | 0.864 | 0.000 | 1.000 | 0.998 |
| HALLMARK_BILE_ACID_METABOLISM | 0.025 | 1.000 | 0.025 | 0.998 | 0.418 | 0.370 | 0.989 | 0.539 | 0.322 | 0.039 | 0.497 | 1.000 | 0.243 | 0.980 | 0.007 | 0.835 | 0.617 |
| HALLMARK_MYOGENESIS | 0.028 | 1.000 | 0.060 | 0.998 | 0.005 | 0.150 | 0.017 | 0.602 | 0.323 | 0.005 | 0.842 | 0.000 | 0.150 | 0.998 | 0.980 | 0.987 | 0.780 |
| HALLMARK_TNFA_SIGNALING_VIA_NFKB | 0.028 | 1.000 | 0.002 | 0.998 | 0.546 | 0.423 | 0.989 | 0.970 | 0.165 | 0.286 | 1.000 | 0.146 | 0.550 | 0.000 | 0.007 | 1.000 | 0.885 |
| HALLMARK_ANGIOGENESIS | 0.028 | 0.039 | 0.161 | 0.998 | 0.625 | 0.578 | 0.669 | 0.165 | 0.331 | 0.933 | 0.858 | 0.853 | 0.741 | 0.252 | 0.050 | 0.945 | 0.998 |
| HALLMARK_REACTIVE_OXIGEN_SPECIES_PATHWAY | 0.092 | 0.017 | 0.799 | 0.998 | 0.307 | 0.578 | 0.989 | 0.748 | 0.820 | 0.029 | 0.114 | 0.145 | 0.178 | 0.705 | 0.808 | 0.600 | 0.998 |
| HALLMARK_P53_PATHWAY | 0.092 | 0.075 | 0.002 | 0.600 | 0.611 | 0.863 | 0.989 | 0.956 | 0.042 | 0.096 | 0.267 | 0.146 | 0.217 | 0.005 | 0.090 | 0.691 | 0.904 |
| HALLMARK_APOPTOSIS | 0.092 | 0.127 | 0.258 | 0.998 | 0.641 | 0.587 | 0.669 | 0.956 | 0.313 | 0.887 | 0.950 | 0.050 | 0.784 | 0.001 | 0.153 | 0.712 | 0.617 |
| HALLMARK_PEROXISOME | 0.096 | 0.771 | 0.253 | 0.650 | 0.191 | 0.225 | 0.862 | 0.539 | 0.050 | 0.153 | 0.842 | 1.000 | 0.894 | 0.980 | 0.007 | 0.382 | 0.060 |
| HALLMARK_ADIPOGENESIS | 0.215 | 0.312 | 0.010 | 0.998 | 0.033 | 0.578 | 0.989 | 0.874 | 0.415 | 0.038 | 0.005 | 0.903 | 0.741 | 0.963 | 0.893 | 0.050 | 0.028 |
| HALLMARK_UV_RESPONSE_UP | 0.215 | 0.730 | 0.247 | 0.998 | 0.542 | 0.863 | 0.571 | 0.970 | 0.043 | 0.324 | 0.158 | 0.140 | 0.666 | 0.000 | 0.461 | 0.714 | 0.050 |
| HALLMARK_ALLOGRAFT_REJECTION | 0.215 | 1.000 | 0.081 | 0.998 | 0.963 | 0.921 | 0.050 | 0.956 | 0.994 | 0.994 | 1.000 | 0.928 | 0.901 | 0.001 | 0.143 | 1.000 | 0.998 |
| HALLMARK_ESTROGEN_RESPONSE_EARLY | 0.239 | 1.000 | 0.019 | 0.998 | 0.160 | 0.379 | 0.989 | 0.539 | 0.030 | 0.025 | 0.001 | 0.853 | 0.195 | 0.416 | 0.194 | 0.792 | 0.617 |
| HALLMARK_EPITHELIAL_MESENCHYMAL_TRANSITION | 0.239 | 0.938 | 0.221 | 0.998 | 0.191 | 0.370 | 0.025 | 0.100 | 0.781 | 0.363 | 0.775 | 0.057 | 0.550 | 0.064 | 0.808 | 0.714 | 0.998 |
| HALLMARK_GLYCOLYSIS | 0.519 | 0.713 | 0.221 | 0.998 | 0.641 | 0.863 | 0.989 | 0.970 | 0.071 | 0.308 | 0.042 | 0.057 | 0.615 | 0.419 | 0.481 | 1.000 | 0.617 |
| HALLMARK_TGF_BETA_SIGNALING | 0.552 | 0.009 | 0.068 | 0.998 | 0.382 | 0.863 | 0.647 | 0.539 | 0.150 | 0.414 | 0.939 | 0.191 | 0.901 | 0.047 | 0.587 | 0.114 | 0.617 |
| HALLMARK_UNFOLDED_PROTEIN_RESPONSE | 0.592 | 0.509 | 0.194 | 0.417 | 0.191 | 0.863 | 0.989 | 0.372 | 0.148 | 0.153 | 0.736 | 0.447 | 0.033 | 0.027 | 0.283 | 0.835 | 0.885 |
| HALLMARK_ANDROGEN_RESPONSE | 0.592 | 0.807 | 0.161 | 0.998 | 0.234 | 0.863 | 0.792 | 0.213 | 0.050 | 0.045 | 0.858 | 0.946 | 0.178 | 0.774 | 0.505 | 1.000 | 0.998 |
| HALLMARK_ESTROGEN_RESPONSE_LATE | 0.592 | 1.000 | 0.068 | 0.998 | 0.307 | 0.488 | 0.757 | 0.727 | 0.133 | 0.153 | 0.291 | 0.769 | 0.550 | 0.047 | 0.125 | 0.714 | 0.617 |
| HALLMARK_PI3K_AKT_MTOR_SIGNALING | 0.592 | 0.039 | 0.221 | 0.998 | 0.415 | 0.863 | 0.488 | 0.182 | 0.600 | 0.933 | 0.012 | 1.000 | 0.550 | 0.548 | 0.127 | 0.194 | 0.773 |
| HALLMARK_COMPLEMENT | 0.592 | 0.938 | 0.081 | 0.998 | 0.546 | 0.863 | 0.333 | 0.165 | 0.940 | 0.784 | 1.000 | 0.140 | 0.550 | 0.162 | 0.011 | 0.835 | 0.617 |
| HALLMARK_DNA_REPAIR | 0.592 | 0.029 | 0.221 | 0.998 | 0.693 | 0.863 | 0.790 | 0.970 | 0.457 | 0.784 | 0.347 | 1.000 | 0.990 | 0.980 | 0.533 | 0.523 | 0.617 |
| HALLMARK_INFLAMMATORY_RESPONSE | 0.592 | 1.000 | 0.019 | 0.998 | 0.846 | 0.863 | 0.905 | 0.956 | 0.994 | 0.860 | 1.000 | 1.000 | 0.901 | 0.000 | 0.125 | 0.940 | 0.998 |
| HALLMARK_APICAL_SURFACE | 0.640 | 1.000 | 0.919 | 0.998 | 0.075 | 0.423 | 0.989 | 0.274 | 0.187 | 0.025 | 0.577 | 0.769 | 0.988 | 0.419 | 0.808 | 0.796 | 0.617 |
| HALLMARK_E2F_TARGETS | 0.721 | 0.275 | 0.221 | 0.998 | 0.167 | 0.863 | 0.989 | 0.970 | 0.090 | 0.887 | 0.888 | 1.000 | 0.033 | 0.963 | 1.000 | 0.791 | 0.998 |
| HALLMARK_G2M_CHECKPOINT | 0.721 | 0.055 | 0.349 | 0.998 | 0.272 | 0.863 | 0.989 | 0.602 | 0.030 | 0.994 | 1.000 | 1.000 | 0.038 | 0.980 | 1.000 | 0.322 | 0.885 |
| HALLMARK_MITOTIC_SPINDLE | 0.721 | 0.000 | 0.194 | 0.998 | 0.388 | 0.863 | 0.083 | 0.045 | 0.135 | 0.994 | 0.903 | 0.908 | 0.741 | 0.710 | 0.808 | 0.050 | 0.009 |
| HALLMARK_MTORC1_SIGNALING | 0.721 | 0.924 | 0.221 | 0.417 | 0.453 | 0.863 | 0.669 | 0.956 | 0.030 | 0.327 | 0.050 | 0.146 | 0.033 | 0.047 | 0.498 | 0.330 | 0.885 |
| HALLMARK_PROTEIN_SECRETION | 0.721 | 0.010 | 0.544 | 0.998 | 0.846 | 0.843 | 0.421 | 0.165 | 0.213 | 0.682 | 0.841 | 0.769 | 0.372 | 0.864 | 0.461 | 0.050 | 0.025 |
| HALLMARK_IL6_JAK_STAT3_SIGNALING | 0.721 | 1.000 | 0.232 | 0.998 | 0.976 | 0.863 | 0.333 | 0.970 | 0.893 | 0.994 | 1.000 | 1.000 | 0.513 | 0.005 | 0.808 | 1.000 | 0.617 |
| HALLMARK_KRAS_SIGNALING_UP | 0.726 | 1.000 | 0.032 | 0.998 | 0.907 | 0.595 | 0.156 | 0.213 | 0.994 | 0.108 | 1.000 | 0.635 | 0.668 | 0.005 | 0.763 | 0.835 | 0.904 |
| HALLMARK_FATTY_ACID_METABOLISM | 0.814 | 1.000 | 0.014 | 0.998 | 0.307 | 0.863 | 0.905 | 0.713 | 0.068 | 0.017 | 0.569 | 1.000 | 0.741 | 0.950 | 0.242 | 0.791 | 0.617 |
| HALLMARK_INTERFERON_GAMMA_RESPONSE | 0.817 | 0.938 | 0.544 | 0.998 | 0.895 | 0.921 | 0.017 | 0.970 | 0.994 | 0.887 | 1.000 | 1.000 | 0.150 | 0.000 | 0.487 | 0.792 | 0.998 |
| HALLMARK_IL2_STAT5_SIGNALING | 0.822 | 1.000 | 0.025 | 0.998 | 0.225 | 0.863 | 0.107 | 0.539 | 0.229 | 0.887 | 0.842 | 0.820 | 0.467 | 0.003 | 0.481 | 0.945 | 0.617 |
| HALLMARK_INTERFERON_ALPHA_RESPONSE | 0.822 | 0.827 | 0.400 | 0.998 | 0.907 | 0.930 | 0.017 | 0.970 | 0.427 | 0.887 | 1.000 | 1.000 | 0.287 | 0.000 | 0.747 | 0.568 | 0.998 |
| HALLMARK_HEME_METABOLISM | 0.874 | 0.564 | 0.994 | 0.998 | 0.542 | 0.863 | 0.905 | 0.713 | 0.030 | 0.731 | 0.615 | 0.057 | 0.666 | 0.804 | 0.808 | 0.835 | 0.882 |
| HALLMARK_APICAL_JUNCTION | 0.935 | 1.000 | 0.572 | 0.998 | 0.307 | 0.488 | 0.443 | 0.165 | 0.307 | 0.005 | 0.558 | 0.733 | 0.741 | 0.682 | 0.481 | 1.000 | 0.317 |

**Figure S4.** Full overview of Molecular Signatures Database (MsigDB) hallmarks that are enriched for caste-dependent differential gene expression. Listed are all pathways that show a significant differential expression signal (FDR < 0.05) in at least one of the tissues or on the cross tissue-level (FDR < 0.1).

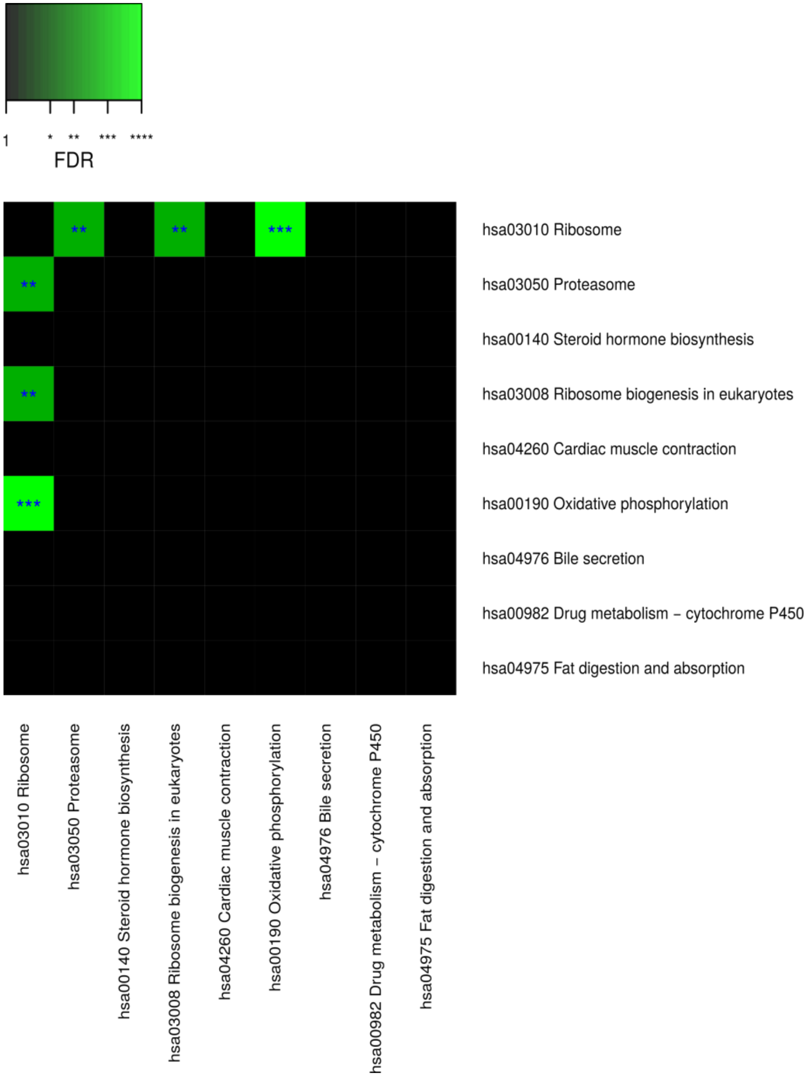


**Figure S5.** Pairwise connectivity of metabolic pathways that are enriched for caste-dependent differential gene expression at the cross-tissue level. Pairwise pathway connectivities were determined based on read counts of all examined 636 samples via a weighted gene co-expression network analysis (WGCNA). Significance was determined against empirically estimated null distributions (see Methods) The following 3 KEGG pathways were excluded since the respective enrichments are likely to be only derivatives from other enrichments (see Results, Figures S7-S9): “hsa05016 Huntington’s disease”,
“hsa05012 Parkinson’s disease”, “hsa050 Alzheimer’s disease”.
*→ FDR < 0.05, ** → FDR < 0.01, *** → FDR < 0.001,
**** → FDR < 0.001 (for exact FDR-values see Results).

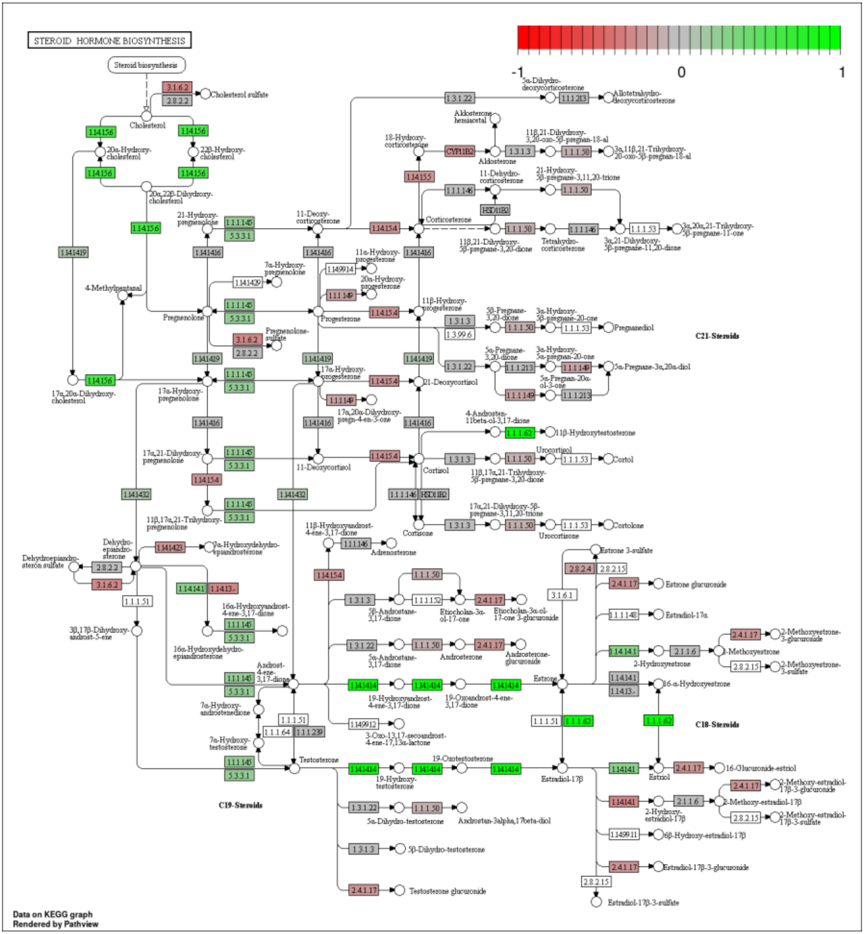


**Figure S6.** Caste-dependent regulation of Kyoto Encyclopedia of Genes and Genomes (KEGG) pathway hsa00140 Steroid hormone biosynthesis at the cross-tissue level. Green/red color depicts the $\log_2$-foldchange (see legend for scale) with direction breeder/non-breeder.

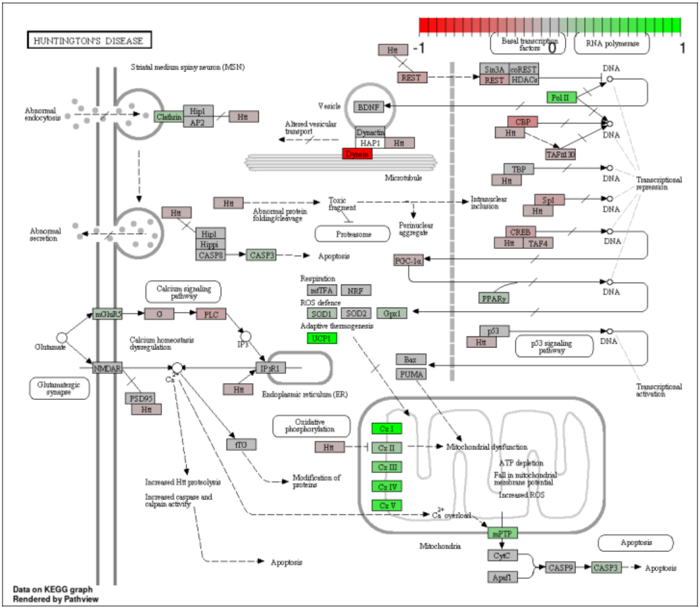


**Figure S7.** Caste-dependent regulation of Kyoto Encyclopedia of Genes and Genomes (KEGG) pathway hsa05016 Huntington's disease at the cross-tissue level. The signal is mainly driven by the differential expression of the mitochondrial respiratory chain. Green/red color depicts the cross-tissue $\log_2$-foldchange (see legend for scale) with direction breeder/non-breeder.

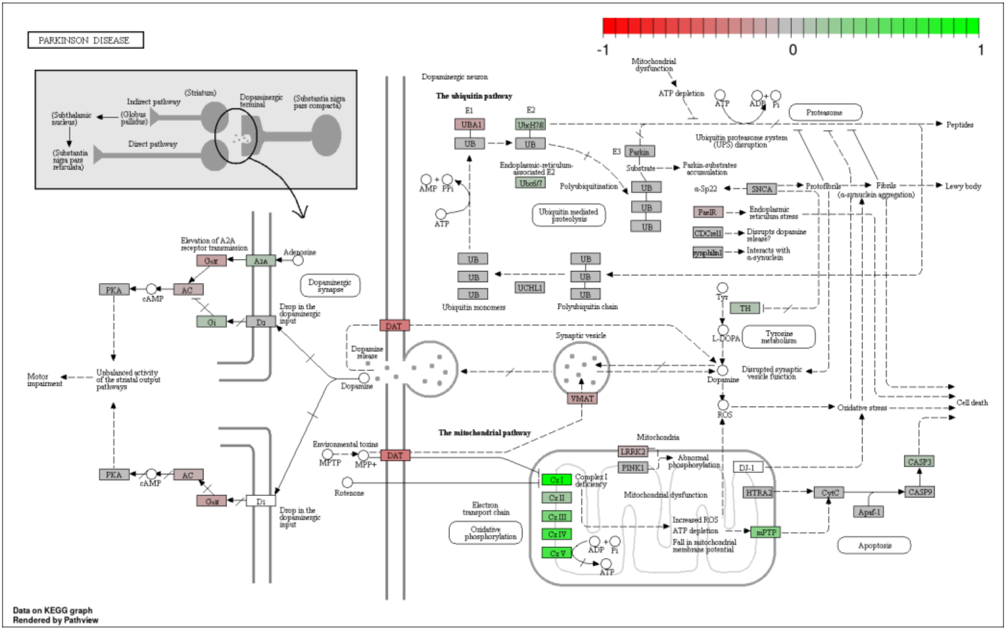


**Figure S8.** Caste-dependent regulation of Kyoto Encyclopedia of Genes and Genomes (KEGG) pathway hsa05012 Parkinson's disease at the cross-tissue level. The signal is mainly driven by the differential expression of the mitochondrial respiratory chain. Green/ red color depicts the cross-tissue $\log_2$-foldchange (see legend for scale) with direction breeder/non-breeder.

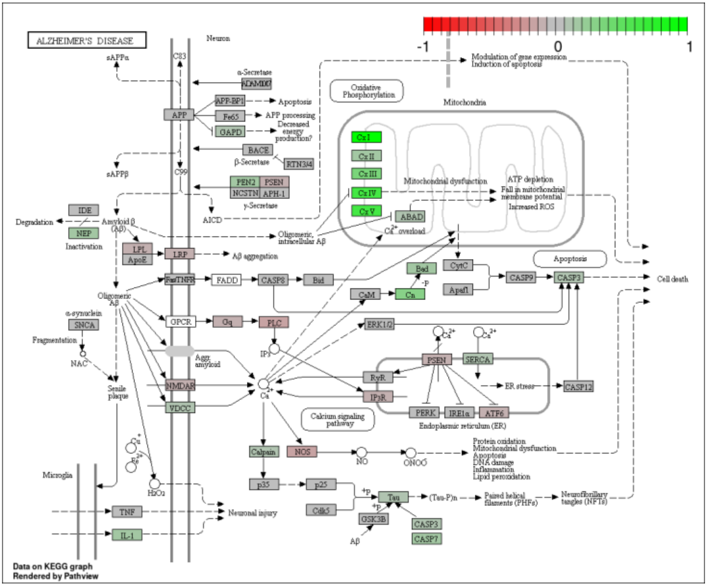


**Figure S9.** Caste-dependent regulation of Kyoto Encyclopedia of Genes and Genomes (KEGG) pathway hsa050160 Alzheimer's disease at the cross-tissue level. The signal is mainly driven by the differential expression of the mitochondrial respiratory chain. Green/ red color depicts the cross-tissue $\log_2$-foldchange (see legend for scale) with direction breeder/non-breeder.
